# Pathological levels of α-synuclein increase basal mitophagy in Parkinson’s dopamine neurons

**DOI:** 10.64898/2026.10.29.762809

**Authors:** Elliot D. Mock, Benjamin Vallin, William McGuinness, Maria Claudia Caiazza, Martha Lavelle, Benjamin Jenkins, Michal Rokicki, Jimena Monzón-Sandoval, Albert Koulman, Caleb Webber, Brent J. Ryan, Richard Wade-Martins

## Abstract

Elevated α-synuclein results in mitochondrial dysfunction, however the exact mechanism is poorly understood. α-Synuclein localises to multiple subcellular sites including mitochondria, where it binds anionic lipids which regulate mitochondrial dynamics through DRP1. We hypothesised that pathogenic levels of α-synuclein lead to dysregulation of mitophagy and lipid-dependent mitochondrial dynamics, causing mitochondrial and neuronal dysfunction. Using induced pluripotent stem cell-dopamine neurons (iPSC-DANs) carrying the α-synuclein gene triplication mutation (*SNCA-Trp*) which causes an early-onset form of Parkinson’s disease (PD), we established that elevated α-synuclein increased basal mitophagy compared to healthy controls and overall loss of mitochondrial mass. This was associated with accumulation of basal mitophagy receptors BNIP3 and NIX and decreased proteasome activity, and was independent of PINK1-Parkin mitophagy. We identified reduced mitochondrial gene expression by single cell RNAseq mirroring mitochondrial depletion. We found increased levels of dynamin-related protein 1 (DRP1)-dependent mitochondrial fission and perturbed levels of anionic lipids, including cardiolipin and phosphatidic acid, two lipids which directly bind to α-synuclein. Pharmacological inhibition of DRP1 reversed the excessive basal mitophagy in *SNCA-Trp* neurons and restored deficits in cellular ATP and calcium release. Finally, we assessed neuronal function and found that impairment of mitochondrial dysfunction by elevated α-synuclein led to reductions in neuronal firing activity. Inhibition of DRP1 rescued the neuronal activity deficit in *SNCA-Trp* neurons, suggesting that relieving – instead of increasing – mitophagy pressure may be beneficial in α-synuclein-driven PD.

## Introduction

α-Synuclein (*SNCA*) accumulation in Lewy bodies and mitochondrial dysfunction are two hallmarks of Parkinson’s disease (PD), which predominantly affects dopamine neurons in the substantia nigra pars compacta.^1^ The α-synuclein A53T gene mutation was the first familial PD mutation to be described, and several other point mutations have followed.^2–3^ Rare *SNCA* duplication or triplication mutations also lead to early-onset PD with dementia with full penetrance, illustrating the pathogenicity caused by elevated α-synuclein levels.^4^ The underlying mechanisms of how α-synuclein drives PD pathogenesis are not yet fully understood. Multiple studies have shown that elevated α-synuclein causes mitochondrial dysfunction in human, mouse and fly models.^5–10^ More specifically, pathological levels of α-synuclein induced mitochondrial fragmentation, loss of complex I activity and mitochondrial membrane potential (MMP) and increased reactive oxygen species (ROS).^5–10^ However, a unifying model as to how α-synuclein elicits these phenotypes and impairs mitochondrial function is lacking.

A primary mechanism of mitochondrial quality control is by mitochondrial degradation or mitophagy. Multiple familial PD genes have been identified that impair mitophagy when mutated, such as *PINK1*, *Parkin*, *LRRK2* and *DJ-1*.^1, 11–12^ In contrast, investigations into the role of α-synuclein on mitophagy showed that α-synuclein mutations mostly increased mitophagy levels.^13–18^ These studies primarily looked into pathogenic point mutations of α-synuclein including A53T and E46K. Determining whether elevated wild-type α-synuclein also modulates mitophagy is particularly relevant because α-synuclein accumulates with age, ageing is the strongest risk factor for PD, and increased α-synuclein burden is a feature of sporadic PD, which accounts for approximately 85% of cases.^19–22^

Although activating mitophagy is generally regarded as beneficial for mitochondrial health by removing damaged mitochondria through activation of PINK1 and Parkin^23^, four recent studies have shown that excessive basal mitophagy mediated by BNIP3 and NIX (or BNIP3L) occurs in people with a rare FBXL4 mutation causing loss of mitochondrial mass.^24–28^ This mutation produces an early infantile onset multi-system disorder called mitochondrial DNA depletion syndrome-13 (MTDPS13), which presents with encephalopathy, hypotonia, lactic acidosis and severe developmental delay. Ganley and colleagues reported that BNIP3/NIX-dependent basal mitophagy is responsible for clearing mitochondria that are still functional, in contrast to PINK1/Parkin mitophagy, which facilitates degradation of damaged mitochondria upon loss of MMP.^29^ Taken together, these studies challenge the view that activating mitophagy is always beneficial and instead suggest that mitochondrial homeostasis depends on appropriately balanced mitophagy.

Several lines of evidence also point to lipid dysregulation in Parkinson’s disease.^30–31^ α-Synuclein binds anionic lipids via its *N*-terminal domain which is essential for targeting to the outer mitochondrial membrane (OMM). Notably, the α-synuclein *N*-terminus, but not the *C*-terminus or NAC domain, functions as a phosphatidic acid (PA) sensor and is essential for mitochondrial fragmentation.^10, 32–33^ It was reported that α-synuclein localizes to mitochondria-ER contact sites (MAMs), which is the principal site for mitochondrial fission orchestrated by the GTPase dynamin-related protein 1 (DRP1). DRP1 activity is regulated by anionic lipids, including cardiolipin (CL) and PA.^34–35^ Once externalised from the inner mitochondrial membrane (IMM) to the OMM, CL increases DRP1 GTPase activity causing mitochondrial fission. In contrast, PA restrains DRP1 activity and is fusogenic.^34, 36^ Whether elevated levels of α-synuclein may offset the balance of mitochondrial fission and fusion mediated by DRP1, CL and PA is unclear.

To model the contribution of increased α-synuclein on mitophagy in a relevant human model system of PD, we have characterised induced pluripotent stem cell (iPSC)-dopamine neurons (iPSC-DANs) from a patient with an *SNCA* gene triplication mutation (*SNCA-Trp*), and compared these to iPSC-DANs from healthy donors. We observed functional PINK1-Parkin-dependent mitophagy, but increased levels of BNIP3-NIX mediated basal mitophagy as a result of downregulated proteasome activity. The increased basal mitophagy led to loss of mitochondrial mass and decreased cellular ATP. Single-cell RNAseq confirmed reduction of mitochondrial and proteasomal gene expression, and dysregulation of lipid-related genes in *SNCA-Trp* iPSC-DANs. Lipidomics revealed several dysregulated lipid classes including cardiolipin and phosphatidic acid, pointing to increased DRP1 activity. Pharmacological inhibition of DRP1 reversed elevated mitophagy and restored ATP, calcium release and neuronal activity to control levels. Taken together, these results provide evidence that elevated α-synuclein pressures mitochondria to undergo mitophagy, putting strain on the remaining pool of functional mitochondria.

## Results

### *SNCA-Trp* iPSC-DANs show elevated basal mitophagy

We differentiated induced pluripotent stem cell (iPSC) lines carrying the Parkinson’s *SNCA* gene triplication (*SNCA-Trp*) mutation and healthy controls into dopamine neurons (iPSC-DANs). Expression of tyrosine hydroxylase (TH) demonstrated a differentiation efficiency into iPSC-DANs of 40-60% with no significant difference between genotypes in accordance with previous reports (Suppl. Fig. 1A,B).^11^ Total α-synuclein immunostaining in TH^+^ iPSC-DANs using the MJFR1 antibody showed a 2-to 3-fold elevation in *SNCA-Trp* iPSC-DANs compared to controls (Suppl. Fig. 2A,B). We confirmed the specificity of the α-synuclein MJFR1 antibody using a *SNCA KO* iPSC-DANs. No elevation of phosphorylated α-synuclein on Ser129 (pS129) was observed in the *SNCA-Trp* iPSC-DANs compared to controls, suggesting that α-synuclein is predominantly in a non-aggregated form (Suppl. Fig. 2C).

To understand how elevated a-synuclein alters mitophagy, we transduced iPSC-DANs on day 20 of the differentiation protocol with a dox-inducible mt-Keima lentiviral vector (Fig. 1A).^11^ Under basal conditions *SNCA-Trp* iPSC-DANs displayed a significant ∼1.5-fold increase of mitophagy index, number and total area of mitolysosomes or red mt-Keima spots (Fig. 1A-D). We confirmed that healthy mitochondria in *SNCA-Trp* iPSC-DANs (represented by the green mt-Keima channel) were more fragmented and smaller in size compared to controls (Fig. 1A,D,E). Importantly, we measured a decrease in total mitochondrial area, which we confirmed by staining for TOM20, suggesting a loss of mitochondrial mass (Fig. 1G, Suppl. Fig. 3A,B). No significant changes were observed for mitochondrial roundness or width to length ratio in the *SNCA-Trp* iPSC-DANs (Suppl. Fig. 3C,D). To investigate the contribution of PINK1/Parkin-mediated mitophagy, we treated neurons with the mitochondrial uncoupler carbonyl cyanide 3-chlorophenylhydrazone (CCCP) and observed 1.5-to 2-fold increase in mitophagy index and 2-to 3-fold increases in mitolysosomes for control and *SNCA-Trp* neurons, respectively (Fig. 1A,H,J). The CCCP-induced mitophagy increase was significantly larger in *SNCA-Trp* iPSC-DANs when normalised to DMSO treated control neurons. When normalised per genotype, however, no significant difference was observed in uncoupler-induced mitophagy, confirming that *SNCA-Trp* iPSC-DANs have a higher basal mitophagy, and that their CCCP-induced mitophagy response is similar to controls (Fig. 1I,K). To investigate this further, we measured phosphorylation of ubiquitin serine-65 (pS65Ub), an established marker of PINK1-Parkin pathway activation.^1, 11^ Few pS65Ub spots were observed under basal conditions in either genotype, while similar strong 3-fold increases were seen upon CCCP treatment (Fig. 1L-N). To have an orthogonal basal mitophagy read-out, we employed the Mtphagy dye^37^ co-localised with Lysotracker Deep Red (Suppl. Fig. 4A,B). We detected increased mitolysosomal spot area in *SNCA-Trp* neurons, which was inhibited by bafilomycin A1 confirming elevated mitophagy flux (Suppl. Fig. 4A,B). These combined results indicate that *SNCA-Trp* iPSC-DANs have elevated basal mitophagy and loss of mitochondrial mass, in the absence of overt changes in the PINK1-Parkin mitophagy pathway.

**Figure 1.**
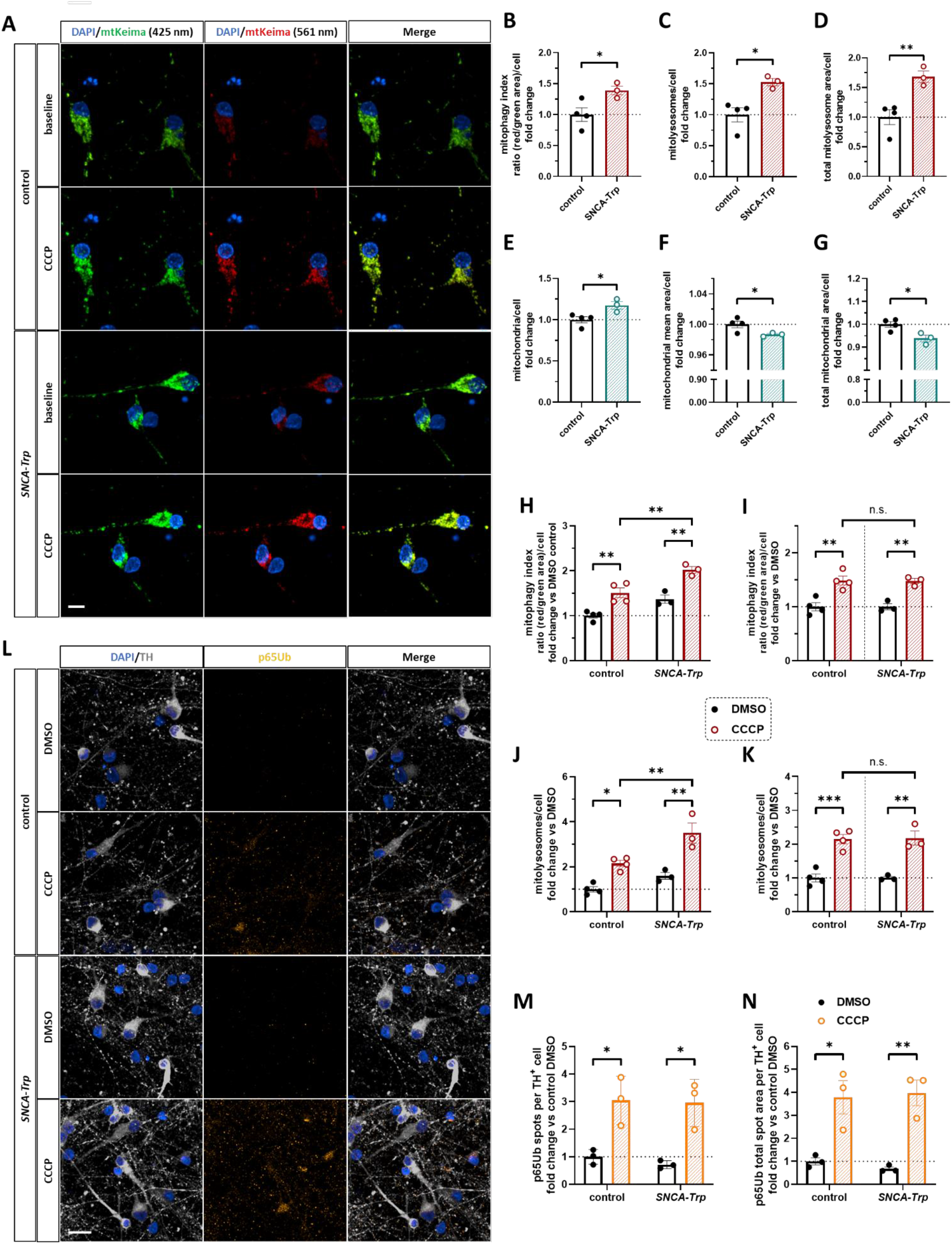
*SNCA-Trp* iPSC-DANs have increased basal mitophagy and mitochondrial loss. **A)** Representative mt-Keima images showing *SNCA-Trp* iPSC-DANs have increased mitophagy compared to controls between D35-50. Cells were treated with doxycycline (1 µg/mL, 48 h). Scale bar = 10 µm. **B-D)** Quantification of **(A)** showing increased mitophagy index **(B)**, mitolysosomes **(C)** and total mitolysosomes area **(D)** in *SNCA-Trp* iPSC-DANs. The mitophagy index was calculated as: (sum of the area of red puncta)/(sum of the area of green puncta) per cell. **E-G)** *SNCA-Trp* iPSC-DANs have an increased number of mitochondria **(D)**, a lower average mitochondrial size **(E)** and lower mitochondrial mass **(G)**. For **B-G)** Data represent the mean ± SEM of 3-4 iPSC lines per genotype averaged over four independent differentiations. *P*-values were calculated using unpaired two-tailed Student’s *t*-test with Welch’s correction. **H+J)** The mitochondrial uncoupler CCCP induced significant increases in mitophagy index **(H)** and mitolysosome puncta **(J)** in the mt-Keima assay. **I+K)** Baseline correction shows similar increases in mitophagy index **(I)** and mitolysosome puncta **(K)** in *SNCA-Trp* and control iPSC-DANs. Cells were treated with doxycycline (1 µg/mL, 48 h), followed by CCCP (5 µM, 4 h). For **H-K)** Data represent the mean ± SEM of 3-4 lines of iPSC-DANs per genotype averaged over four independent differentiations. *P*-values were calculated using two-way ANOVA with Tukey’s multiple comparisons test. **L)** Representative images of pS65Ub staining in TH-positive iPSC-DANs showing low basal expression and induction with CCCP treatment (10 µM, 6 h) in both genotypes at D35-50. Scale bar = 20 µm. **M,N)** Quantification of pS65Ub induction by CCCP showing increases in spot number **(M)** and total spot area **(N)** per TH^+^ cell. For **M,N)** Data represent the mean ± SEM of 3-4 lines of iPSC-DANs per genotype averaged over three independent differentiations. *P*-values were calculated using two-way ANOVA with Tukey’s multiple comparisons test. *, *P* < 0.05; **, *P* < 0.01; ***, *P* < 0.001.

### BNIP3 and NIX are upregulated in *SNCA-Trp* iPSC-DANs, while proteasomal activity is impaired

To understand the apparent increase in basal mitophagy in the absence of overt PINK1 activation, the levels of PINK1-independent mitophagy mediators were assessed. BNIP3 and NIX are established markers of receptor-mediated basal mitophagy and are anchored on the OMM where they can directly recruit the autophagy machinery.^38^ We measured significantly increased protein expression for BNIP3 and NIX in *SNCA-Trp* TH^+^ iPSC-DANs (Fig. 2A-C). Given BNIP3 and NIX are regulated by FBXL4-mediated proteasomal degradation,^24–28, 39–40^ we assessed whether proteasomal activity was altered in *SNCA-Trp* neurons and if this is sufficient to modulate BNIP3/NIX and neuronal mitophagy. We observed a 20% reduction of chymotrypsin-like proteasome activity in *SNCA-Trp* iPSC-DANs (Fig. 2D,E). We reasoned that pharmacological proteasome inhibition may enhance mitophagy through BNIP3 and NIX accumulation similar to *SNCA* triplication. We performed a concentration-response experiment using the proteasome inhibitor epoxomicin in healthy control (IC_50_ = 0.93 nM, 95% CI: 0.23 – 1.7 nM) and *SNCA-Trp* iPSC-DANs (IC_50_ = 0.90 nM, 95% CI: 0.25 – 1.6 nM) (Suppl. Fig. 5). Next, we confirmed that epoxomicin (5 nM, 48 h) increases mitophagy in the mt-Keima assay in control and *SNCA-Trp* iPSC-DANs (Fig. 2F,G). This was accompanied by a significant elevation of BNIP3, but not NIX expression in TH^+^ cells (Fig. 2H-J). Similar to proteasome inhibition, the E1 neddylation inhibitor MLN4924, which blocks Cullin-RING E3 ligase activity, significantly enhanced BNIP3 and NIX levels and mitophagy in the mt-Keima assay (Fig. 2F-J). Taken together, these results indicate that BNIP3 and NIX play an important role in the basal mitophagy pathway in dopamine neurons and this pathway is elevated in *SNCA-Trp* iPSC-DANs, in part through inhibition of their constitutive degradation via the proteasome.

**Figure 2.**
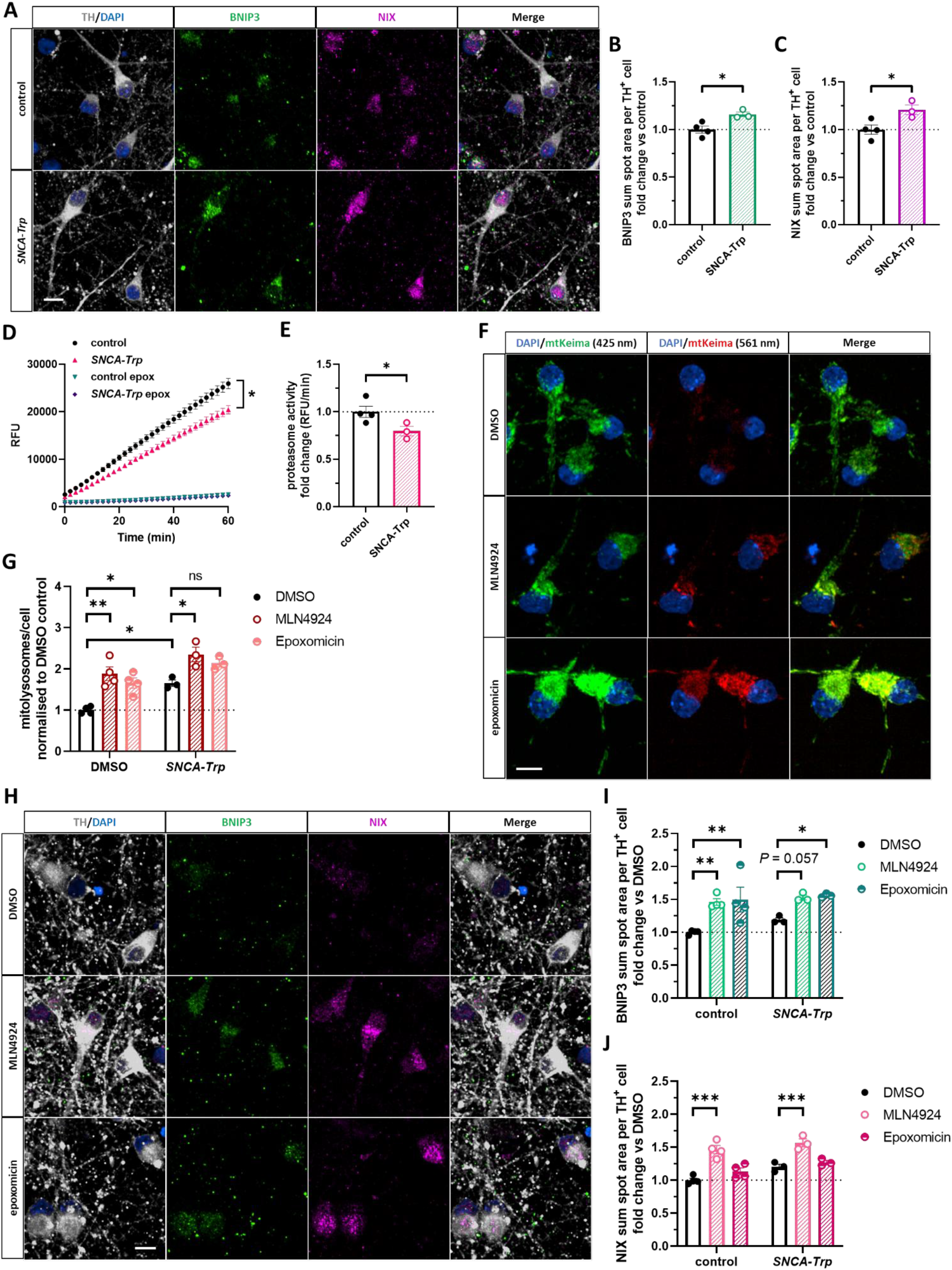
*SNCA-Trp* iPSC-DANs have elevated NIX and BNIP3 expression and reduced proteasome activity. **A)** Representative images showing increased BNIP3 and NIX expression in *SNCA-Trp* iPSC-DANs versus controls. **B,C)** Quantification of **(A)** for BNIP3 **(B)** and NIX **(C)**. **D)** Chymotrypsin-like (β5) proteasome activity measured using the turnover of the fluorogenic substrate Suc-LLVY-AMC is significantly decreased in *SNCA-Trp* iPSC-DANs. **E**) Quantification of the turnover rate (RFU/min). For **B-E**) Data represent the mean ± SEM of 3-4 lines of iPSC-DANs per genotype averaged for three independent differentiations. *P*-values were calculated using unpaired, two-tailed Student’s *t*-test with Welch’s correction. **F)** Representative mt-Keima images of *SNCA-Trp* iPSC-DANs showing increased mitophagy after blocking E1 neddylation (MLN4924, 1 µM, 48 h) or proteasome (epoxomicin, 5 nM, 48 h) activity. Cells were co-treated with doxycycline (1 µg/mL, 48 h). **G)** mt-Keima quantification of mitolysosomes related to **(F)**. Data represent the mean ± SEM of 3-4 lines of iPSC-DANs per genotype averaged for three independent differentiations. *P*-values were calculated using two-way ANOVA with Tukey’s multiple comparisons test. **H)** Representative images showing increased BNIP3 and NIX expression after treatment with MLN4924 (1 µM, 48 h) or epoxomicin (5 nM, 48 h) in *SNCA-Trp* TH^+^ iPSC-DANs. **I-J)** Immunocytochemistry quantification of **(H)** showing increased BNIP3 **(I)** and NIX **(J)** total spot area after treating with MLN4924 (1 µM, 48 h) or epoxomicin (5 nM, 48 h). For **I-J**) Data represent the mean ± SEM of 3-4 lines of iPSC-DANs per genotype averaged for three independent differentiations. *P*-values were calculated using two-way ANOVA with Tukey’s multiple comparisons test. *, *P* < 0.05; **, *P* < 0.01; ***, *P* < 0.001. Scale bar = 10 µm for **A**, **F**, **H**.

### scRNAseq reveals mitochondrial, proteasome and lipid genes downregulated in *SNCA-Trp* iPSC-DANs

To investigate the transcriptional changes that underpin the observed mitophagy phenotype in iPSC-DANs carrying the *SNCA-Trp* mutation, we performed scRNA-seq on *SNCA-Trp* and healthy control iPSC-DANs. 1,132 single cells were deeply sequenced using SMART-Seq technology to obtain full-length transcriptomic profiles of iPSC-DANs of 3 healthy control and 3 *SNCA-Trp* lines. Three main clusters were identified including ‘neurons’, ‘astrocytes’, and ‘fibroblast-like’, as observed by the expression of neuronal marker SNAP25, astrocytic marker AQP4, or fibroblast marker COL1A1 (Fig. 3A-E). Broad TH expression was observed in the neuronal cluster (Fig. 3C).

**Figure 3.**
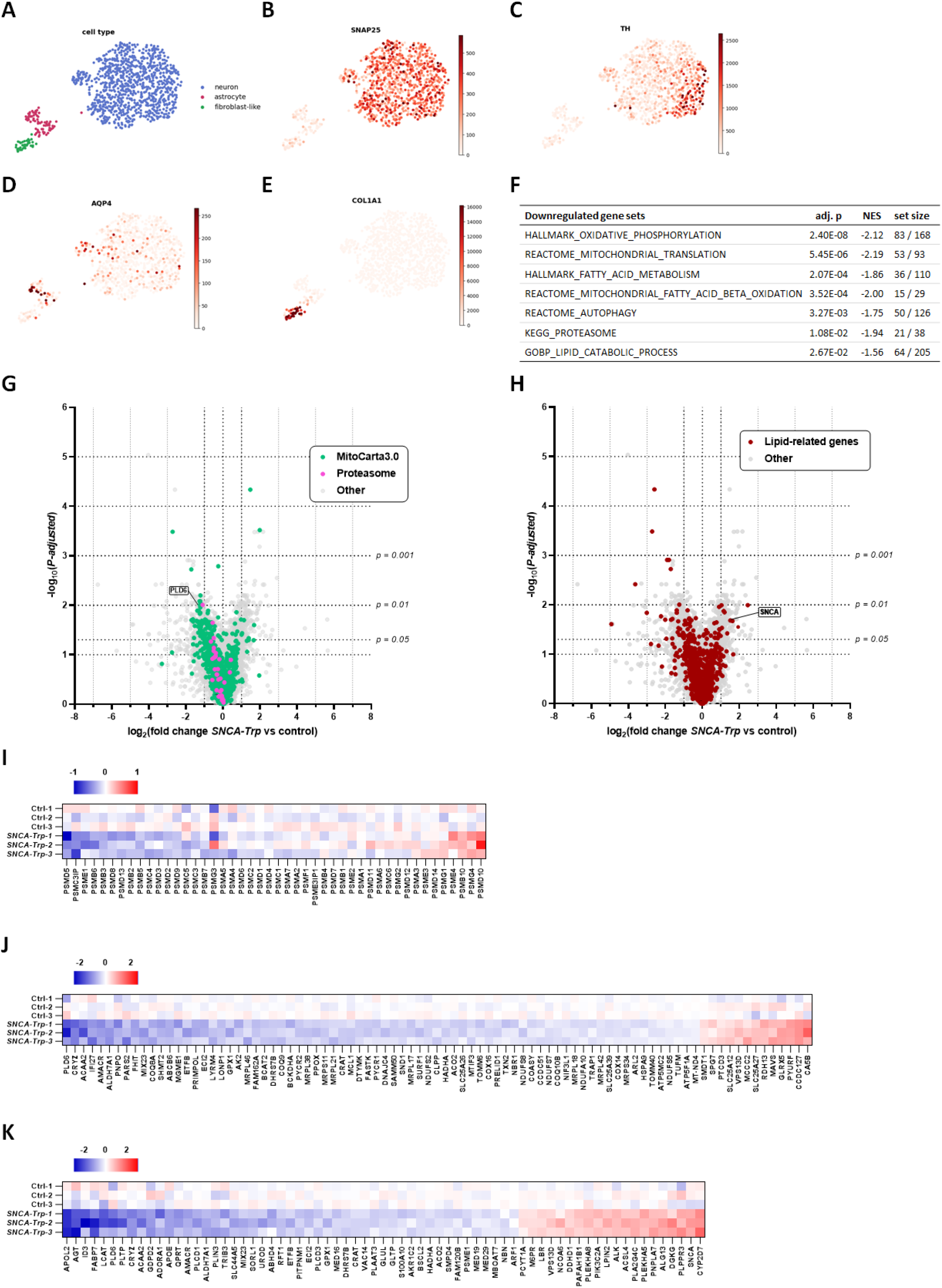
scRNA-seq reveals downregulated expression of mitochondrial, proteasomal and lipid-related genes in *SNCA-Trp* iPSC-DANs. **A-E**) Scatter plot in UMAP basis single cells coloured by cell type (**A**), pan-neuronal marker SNAP25 (**B**), dopamine neuron marker TH (**C**), astrocytic marker AQP4 (**D**), and fibroblast-like marker COL1A1 (**E**). **F**) Table summarizing GSEA results for a selection of gene sets negatively enriched in *SNCA-Trp* vs healthy control iPSC-DANs. adj. p: adjusted *P*-value; NES: normalized enrichment score; set size: number of genes contributing to significant enrichment over total number of genes in gene set. **G)** Volcano plot depicting downregulated expression of mitochondrial (MitoCarta3.0, green) and proteasome subunit genes (magenta) from the DGEA. **H)** Volcano plot showing differentially expressed lipid-related genes, including elevated *SNCA* from the DGEA. **I-K**) Heat maps displaying log_2_ transformed differential expression per line of iPSC-DANs for all proteasome subunits (**I**), and for significantly (*P*-adjusted < 0.05) differentially expressed mitochondrial genes (**J**), and lipid-related genes (**K**). For **G,H**), data represent the mean log_2_ fold change ratio from the DGEA for 3 controls and 3 *SNCA-Trp* lines of iPSC-DANs with Benjamini-Hochberg adjusted *P*-value (FDR < 0.05).

Differential gene expression analysis (DGEA) was performed on the neuron cluster of 952 single cells with an average of 158 cells per line with 12,096 detected total genes after excluding non-protein-coding and lowly expressed genes. 1024 differentially expressed genes at *P*-adjusted threshold = 0.05 were observed in *SNCA-Trp* iPSC-DANs compared to control (Suppl. Table 1). Gene set enrichment analysis (GSEA) provided gene sets negatively enriched in *SNCA-Trp* iPSC-DANs including mitochondrial genes related to oxidative phosphorylation, translation and fatty acid beta oxidation, autophagy, proteasome subunits and lipid-related genes (Fig. 3F-H, Suppl. Table 2). The significant downregulation of proteasome subunits in *SNCA-Trp* iPSC-DANs lends support to the observed loss of proteasome activity in the *SNCA-Trp* neurons (Fig. 3F,G,I). Mitochondrial-related genes were identified using the MitoCarta3.0 database^39^, showing a significant decrease in gene expression for electron transport chain (ETC) subunits of complexes I, IV, and V, and genes involved in the citric acid cycle, fatty acid beta oxidation and the TOM complex (Fig. 3F,G,J). We also observed differential expression of autophagy genes corresponding to impaired autophagic flux as we previously reported for *SNCA-Trp* iPSC-DANs (Fig. 3F).^40^ This included reduced expression of *MAP1LC3B* and *NBR1* (Suppl. Fig. 6A,B). Lastly, broad expression changes were observed in lipid-related gene sets including lipid and fatty acid metabolism and cholesterol homeostasis (Fig. 3F,H). Reassuringly, *SNCA* expression was robustly increased by two-fold in *SNCA-Trp* iPSC-DANs (Fig. 3H,K).

### *SNCA-Trp* iPSC-DANs display lipid dysregulation and elevated cardiolipin

To assess the consequences of differential gene expression of lipid-related genes in the *SNCA-Trp* iPSC-DANs and given the established role of α-synuclein in lipid biology^41^, we performed global lipidomic profiling. 494 individual lipid species across 33 lipid classes were measured (Suppl. Table 3). We observed significant increases of cardiolipin (CL) lipid species and several plasmalogen-type phosphatidylcholine lipids (PC-Ps) (Fig. 4A). Decreased polyunsaturated fatty acids (PUFAs) were detected, including arachidonic (AA, 20:4) and docosahexaenoic acid (DHA, 22:6) (Fig. 4A,B). We noted increased gene expression of acyl-coenzyme A (CoA) synthetase long-chain family member 4 (ACSL4) in *SNCA-Trp* iPSC-DANs (Fig. 3G,J), which prefers PUFAs such as AA as substrate and may increase sensitivity to ferroptosis.^42–43^ Externalisation of CL from the inner to the outer mitochondrial membrane (OMM) – where it is recognized by the autophagy machinery – is an established mitophagy pathway.^44^ Interestingly, we observed decreased gene expression of phospholipase PLD6 (mitoPLD) (Fig. 3F,I), which metabolises CL to phosphatidic acid (PA) on the OMM and thereby regulates mitochondrial fission and mitophagy through modulation of DRP1 activity (Fig. 5A).^34, 36^ We detected an increase of total CL in *SNCA-Trp* iPSC-DANs, whereas PA species were decreased, pointing to a potential second mechanism of increased mitophagy in *SNCA-Trp* iPSC-DANs (Fig. 4C,D).

**Figure 4.**
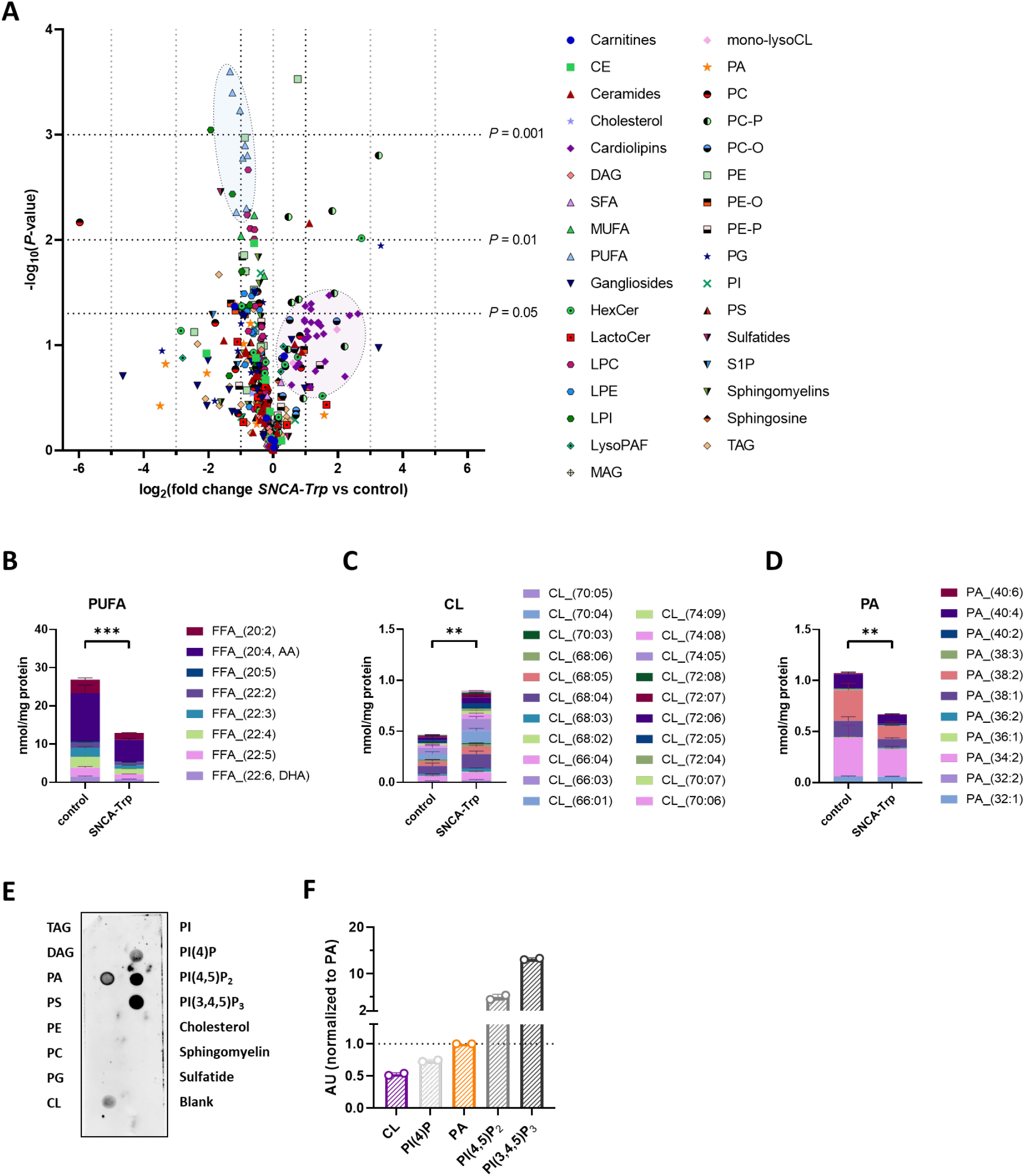
Lipid dysregulation in *SNCA-Trp* iPSC-DANs and lipid preference of α-synuclein. **A)** Volcano plot depicting all lipid species detected. *P*-values were calculated using Student’s *t*-test (unpaired, two-tailed) with Benjamini–Hochberg correction (10% FDR). Abbreviations: CE = cholesterylester; DAG = diacylglycerol; SFA = saturated fatty acid; MUFA = monounsaturated fatty acid; PUFA = polyunsaturated fatty acid; HexCer = hexosylceramide; LactoCer = lactosylceramide; LPC = lysophosphatidylcholine; LPE = lysophosphatidylethanolamine; LPI = lysophosphatidylinositide; LysoPAF = lysoplatelet-activating factor; MAG = monoacylglycerol; mono-lysoCL = mono-lysocardiolipin; CL = cardiolipin; PA = phosphatidic acid; PC = phosphatidylcholine; PC-O = alkyl phosphatidylcholine; PC-P = plasmenyl phosphatidylcholine; PE = phosphatidylethanolamine; PE-O = alkyl phosphatidylethanolamine; PE-P = plasmenyl phosphatidylethanolamine; PG = phosphatidylglycerol; PI = phosphatidylinositide; PS = phoshatidylserine; S1P = sphingosine-1-phosphate; TAG = triacylglycerol. **B-D)** Sum of all (**B**) PUFA, (**C**) CL, and (**D**) PA lipid species detected. For **A-D**) Data represent mean ± SEM from 3 lines of iPSC-DANs per genotype each with three technical replicates from 1 differentiation. *P*-values were calculated using two-way ANOVA with Tukey’s multiple comparisons test. **E)** Representative dot blot showing affinity of human recombinant full length α-synuclein for anionic lipids. **F**) Quantification of dot blot in **(E).** Data ± SEM for two independent replicates. **, *P* < 0.01.

**Figure 5.**
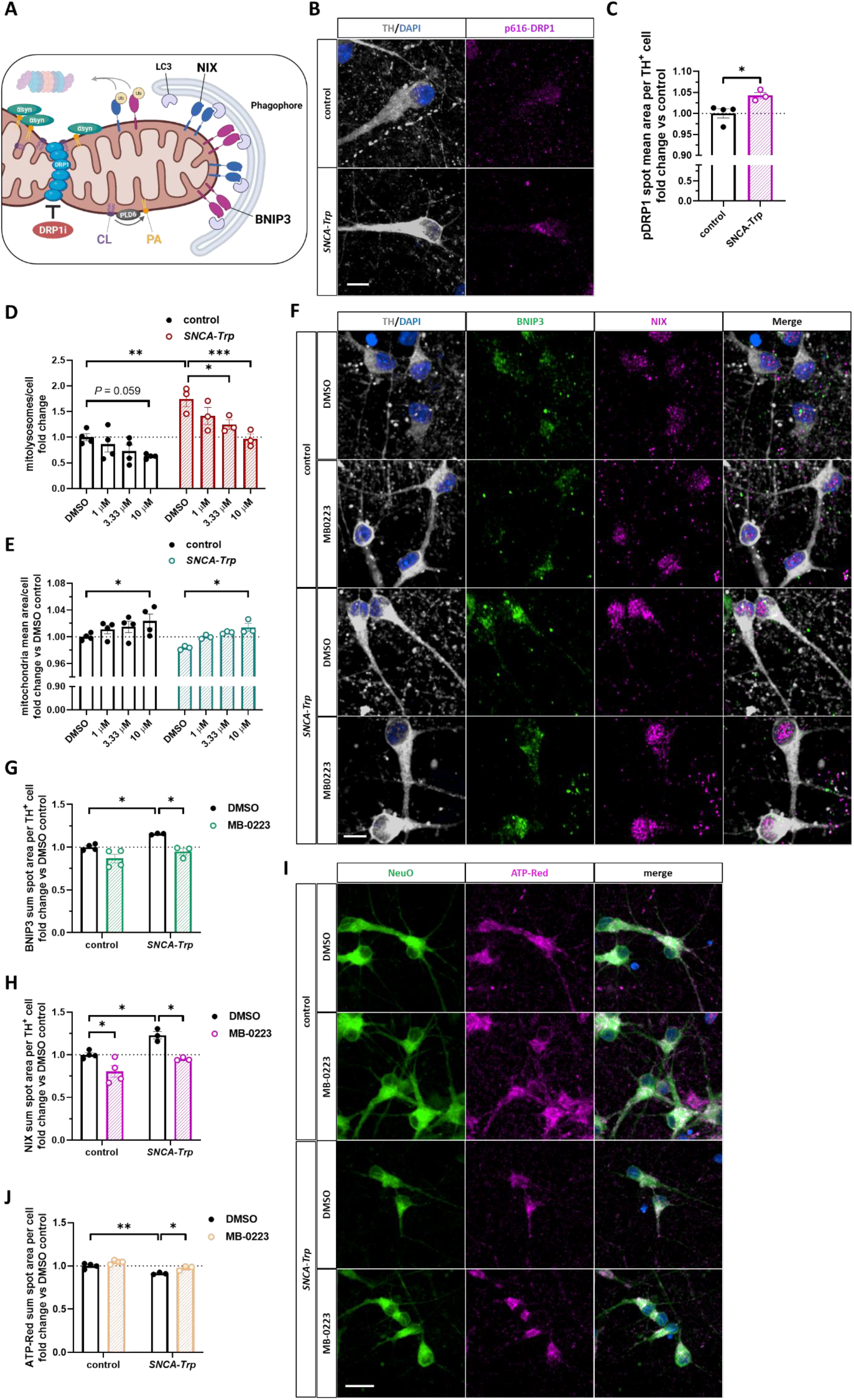
DRP1 inhibition restores mitophagy and mitochondrial phenotypes in *SNCA-Trp* iPSC-DANs. **A)** Proposed model depicting how DRP1 inhibition may reverse elevated mitophagy in *SNCA-Trp* iPSC-DANs. **B)** Representative images showing increased pSer616-DRP1 puncta in *SNCA-Trp* iPSC-DANs compared to controls. Scale bar = 10 µm. **C)** Quantification of p616-DRP1 immunofluorescence. Data represent the mean ± SEM of 3-4 lines of iPSC-DANs per genotype averaged for three independent differentiations. *P* values were calculated unpaired, two-tailed Student’s *t*-test with Welch’s correction. **D)** mt-Keima quantification showing dose-dependent reduction of mitolysosomes and **E)** increase in mitochondrial mean area by treatment with MB-0223 (48 h). Cells were cotreated with doxycycline (1 µg/mL, 48 h). **F)** Representative images showing increased BNIP3 and NIX expression in *SNCA-Trp* dopaminergic neurons and rescue with MB-0223 (10 µM, 48 h). Scale bar = 10 µm. **G,H)** Immunocytochemistry quantification showing reduced expression of BNIP3 **(G)** and NIX **(H)** in TH^+^ iPSC-DANs after treating with MB-0223 (3 µM, 48 h). For **D,E,G,H**) Data represent the mean ± SEM of 3-4 lines of iPSC-DANs per genotype averaged for three independent differentiations. **I)** Representative images of NeuO^+^ neurons stained with ATP sensitive dye BiotrackerATP-Red. Scale bar = 20 µm. **J**) Quantification of BiotrackerATP-Red sum spot area. Data represent the mean ± SEM of 3-4 lines of iPSC-DANs per genotype averaged for four independent differentiations. *P* values were calculated using two-way ANOVA with Tukey’s multiple comparisons test. *, *P* < 0.05; **, *P* < 0.01; ***, *P* < 0.001.

Because α-synuclein is well-established to bind anionic lipids such as CL and PA with its basic *N*-terminal domain, we assessed whether alpha-synuclein has a specific lipid binding preference, which can affect downstream DRP1 activity.^30, 32, 45^ Using a dot blot containing neutral, zwitterionic and anionic lipids we evaluated their specific interaction with recombinant full length human α-synuclein (Fig. 4E). α-Synuclein showed affinity for the anionic lipids PA, CL, phosphatidylinositol-4-phosphate (PI(4)P), phosphatidylinositol-4,5-diphosphate (PI(4,5)P_2_) and phosphatidylinositol-3,4,5-triphosphate (PI(4,5)P_3_). Remarkably, no immunoreactivity was observed for the anionic phospholipids phosphatidylserine (PS), phosphatidylglycerol (PG), and PI, or the sulfolipid sulfatide, suggesting that α-synuclein displays lipid binding specificity. Indeed, quantification showed that α-synuclein preferred PA over CL by 2-fold, although much stronger affinity was apparent for PI(4,5)P_2_ and PI(3,4,5)P_3_ (Fig. 4F). Taken together, these data suggest that dysregulated mitochondrial lipid homeostasis induced by elevated alpha-synuclein may be mediated by its specific anionic lipid preference.

### DRP1 inhibition reverses mitochondrial phenotypes in *SNCA-Trp* iPSC-DANs

Given the involvement of DRP1 in mitophagy and the regulation of its activity by CL and PA levels^35, 46^, we reasoned that DRP1 inhibition may reverse the observed mitochondrial phenotypes. In particular, we hypothesised that in the presence of excessive amounts of α-synuclein, DRP1 may become over-activated due increased levels of CL and sequestration of PA by α-synuclein on the OMM (Fig. 5A). In agreement, we observed increased p616-DRP1 immunofluorescence, considered to be a marker of DRP1 fission activity^47^, in *SNCA-Trp* TH^+^ iPSC-DANs compared to controls (Fig. 5B-C). This was consistent with our previous work showing elevated p616-DRP1 in the midbrain of SNCA-OVX mice.^6^ Here, we found that treatment with the DRP1 inhibitor MB-0223^48^ was able to downregulate mitophagy in a dose-dependent manner in the mt-Keima assay in both *SNCA-Trp* and control iPSC-DANs (Fig. 5D). We observed a concomitant dose-dependent elevation of mitochondrial size, suggesting MB-0223 increases mitochondrial fusion (Fig. 5E). This was confirmed using immunofluorescence of the mitochondrial marker TOM20 in TH^+^ iPSC-DANs (Suppl. Fig. 7A,B). MB-0223 also reduced expression of BNIP3 and NIX in the *SNCA-Trp* neurons to control levels (Fig. 5F-H, Suppl. Fig. 7). Furthermore, *SNCA-Trp* iPSC-DANs showed significantly reduced ATP levels using the cellular ATP sensor BiotrackerATP-Red^49^, and was reversed by DRP1 blockade (Fig. 5I, J). We corroborated the findings on ATP production using an orthogonal luminescence assay read-out (Suppl. Fig. 7C).

### Impaired synaptic activity in *SNCA-Trp* iPSC-DANs can be rescued by inhibition of DRP1

Mitochondrial function and calcium dynamics govern neuronal and synaptic activity.^50^ Previously, we reported that *SNCA-Trp* iPSC-DANs display a deficit in ionomycin-induced Ca^2+^-release using the Fura2-AM assay.^51^ We reasoned that inhibition of DRP1 may restore mitochondrial Ca^2+^-buffering capacity, which may improve neuronal health. Indeed, MB-0223 rescued the impaired Ca^2+^-release in *SNCA-Trp* iPSC-DANs (Fig. 6A,B). To assess neuronal and synaptic activity in the iPSC-DANs, we set up a multi-electrode array (MEA) assay. We optimised conditions for maximal neuronal activity by replacing Neurobasal for BrainPhys media and by adding rat astrocytes to support neuronal maturation and decrease variability (Suppl. Fig. 8A-G). We observed pacemaking activity and bursting activity, with neurons firing in synchronised network bursts starting from 1-2 weeks after replating on D20 in both *SNCA-Trp* and control iPSC-DANs (Suppl. Fig. 9H-I). CNQX and D-AP5 abolished all network firing (Suppl. Fig. 9A-C), confirming that network firing is dependent on AMPA and NMDA receptors as previously reported.^52^ In our scRNAseq dataset we observed that 60-90% of all cells expressed *SLC17A6* (VGLUT2), but not the cortical marker *SLC17A7* (VGLUT1). We confirmed that TH^+^ iPSC-DANs co-expressed VGLUT2 in the soma and axons (Suppl. Fig. 10), indicating that these dopamine neurons may release dopamine and glutamate. Previously, we reported that control and *SNCA-Trp* iPSC-DANs have similar glutamate release events and levels of VGLUT2 as measured by immunofluorescence.^53^ Next, we evaluated neuronal activity longitudinally in *SNCA-Trp* and control iPSC-DANs. After 50 days of maturation (D60) *SNCA-Trp* iPSC-DANs displayed a plateau in their neuronal firing and bursting rates, which was significantly lower compared to controls (Fig. 6C,D). This was also apparent in the number of network bursts, although this metric was more variable overall (Fig. 6E). Importantly, no difference was seen in the number of active electrodes, suggesting an intrinsic neuronal activity deficit in *SNCA-Trp* iPSC-DANs (Fig. 6F). Random spike activity with more spiking outside network bursts was observed in *SNCA-Trp* iPSC-DANs (Fig. 6G-H).

**Figure 6.**
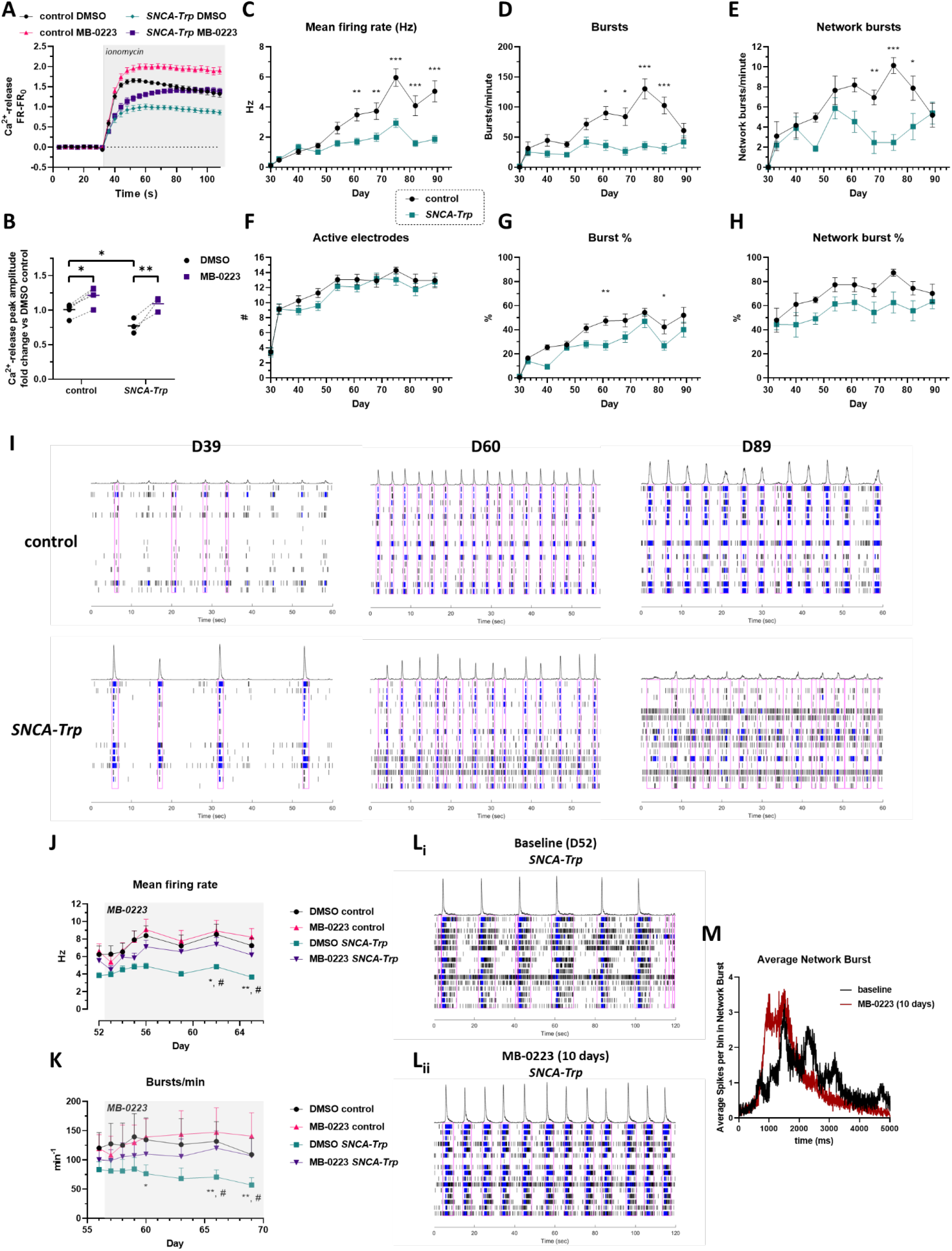
Impaired neuronal activity in *SNCA-Trp* iPSC-DANs is reversed with DRP1 inhibitor MB-0223. **A)** Fura2-AM response traces of ionomycin-induced Ca^2+^-release showing that MB-0223 (3 µM, 48 h) restores the release deficit in *SNCA-Trp* iPSC-DANs. **B)** Quantification of peak amplitude in the Fura2 assay. For **A,B)** Data represent the mean ± SEM of 3-4 lines of iPSC-DANs per genotype averaged for two independent differentiations. *P*-values were calculated using two-way ANOVA with Tukey’s multiple comparisons test. *, *P* < 0.05; **, *P* < 0.01; ***, *P* < 0.001. **C-H**) Longitudinal evaluation of neuronal activity in control and *SNCA-Trp* iPSC-DANs from D30-D90 showing mean firing rate in Hz **(C)**, bursts/minute **(D)**, network bursts/minute **(E)**, # active electrodes **(F)**, burst % **(G)**, and network burst % **(H)**. For **C-H)** Data represent the mean ± SEM average of 3 lines of iPSC-DANs per genotype averaged for three independent differentiations. *P*-values were calculated using two-way ANOVA with Tukey’s multiple comparisons test. **I**) Representative spike raster plots of MEA recordings at three different timepoints (D39, D60, D89) showing development of synchronised network firing in control and *SNCA-Trp* iPSC-DANs. Single spikes are indicated in black, bursts in blue and network bursts are highlighted with pink boxes. **J,K)** Continuous treatment with MB-0223 (3 µM) starting from D52 restores **(J)** mean firing activity and **(K)** bursting activity in *SNCA-Trp* iPSC-DANs. **Li+Lii)** Representative spike raster plots showing the increased firing and more synchronised network activity after treating with MB-0223 (3 µM) for 10 days. **M)** Average network bursting trace showing a more regular shape after treatment with MB-0223. For **J-K)** Data represent mean ± SEM of 3 lines of iPSC-DANs per genotype for two differentiations. *P*-values were calculated using two-way ANOVA with Tukey’s multiple comparisons test. *, *P* < 0.05; **, *P* < 0.01; *** *P* < 0.001 for DMSO *SNCA-Trp* vs MB-0223 *SNCA-Trp*; ^#^, *P* < 0.05 for DMSO control vs. DMSO *SNCA-Trp*.

We hypothesised that reducing the excessive mitophagy by DRP1 inhibition may alleviate the neuronal activity deficit in *SNCA-Trp* iPSC-DANs. We found that continuous treatment with MB-0223 for two weeks starting from 42 days of maturation (D52), reversed the impaired neuronal firing and bursting activity (Fig. 6J,K)). More synchronised network bursting activity was observed after 10 days, showing a more regular shape of the average of all network bursts (Fig. 6L,M). Taken together, these data indicate impairment of neuronal function in *SNCA-Trp* iPSC-DANs compared to controls, which may be rescued by lessening mitophagy through DRP1 blockade.

## Discussion

In our work, we use iPSC-DANs from a PD patient harbouring the *SNCA-Trp* mutation and healthy controls to show that elevated levels of α-synuclein lead to increased basal mitophagy. Specifically, elevated α-synuclein increases mitophagy initiation through concerted upregulation of BNIP3, NIX and cardiolipin. In contrast, PINK1/Parkin-mediated mitophagy, which is activated upon loss of mitochondrial membrane potential, was not altered in *SNCA-Trp* neurons. Collectively, our results provide a novel mechanism as to how α-synuclein leads to mitochondrial impairment and proposes increased basal mitophagy initiation and loss of mitochondrial mass as a potential driver of PD pathology.

Although α-synuclein and mitochondrial dysfunction are each implicated in the development of PD, a unified mechanism as to how α-synuclein induces mitochondrial damage has remained elusive. It is widely established that α-synuclein expression increases with age, is elevated in PD patients, forms the main component of Lewy bodies and is the strongest genetic risk factor for developing PD by GWAS.^19–21^ Notably, aging is the highest risk factor for developing PD, which suggests that age-dependent α-synuclein accumulation may be one of its underlying mechanisms.^22^ Importantly, duplication and triplication mutations of the *SNCA* gene lead to early-onset PD with dementia, indicating that increased levels of wild-type α-synuclein are sufficient to induce PD pathology.^4^

Previous studies have highlighted a role for α-synuclein in mitophagy, showing predominantly increased mitophagy by cardiolipin externalisation.^13–18^ For the most part, these reports investigated familial mutations of α-synuclein including A53T and E46K, or aggregated α-synuclein, and did not evaluate the effect of increased wild-type α-synuclein as we have in this study. Recently, Vanderberghe and co-workers reported impaired mitophagy flux in iPSC-derived neurons with elevated α-synuclein.^54^ Importantly, this was only observed when valinomycin was used as a mitophagy inducer and did not occur under basal conditions. In contrast, we have not observed impairments in mitophagy induction in *SNCA-Trp* iPSC-DANs upon treatment with CCCP using mt-Keima or pSer65Ub immunofluorescence in TH^+^ -neurons (Fig. 1).

We and others have previously shown that elevated α-synuclein impairs lysosomal function and hydrolase activity in the absence of overt aggregation.^40, 55–57^ One could envision that the combined actions elicited by α-synuclein of increased mitophagy initiation and reduced lysosomal degradation may lead to traffic jams of damaged mitochondria. Moreover, α-synuclein was reported to stall mitolysosomes via an interaction with Miro1.^58^ These combined effects of α-synuclein may provide a plausible mechanism for the emergence of Lewy-body-like structures which incorporate mitochondria as well as other organelles.^59–61^ Of note, two recent studies indicated that mitophagy is increased with age in mouse brain and substantia nigra, while autophagy was not affected.^62–63^ In line with those findings, we previously reported depletion of mitochondrial genes in dopaminergic neurons from aged mice, while BNIP3 was increased.^51^

Our current study puts forward receptor-mediated mitophagy via BNIP3 and NIX as a novel mechanism by which α-synuclein can induce basal mitophagy. In *SNCA-Trp* iPSC-DANs we found elevated protein levels of NIX and BNIP3 under basal conditions (Fig. 2). Recently, it was shown that loss-of-function mutations in FBXL4, which is part of the mitochondrial SCF-FBXL4 ubiquitin E3 ligase complex and constitutively ubiquitinates BNIP3 and NIX, lead to excessive mitophagy and caused the mitochondrial disease MTDPS13.^24–26, 64–65^ The described cases died at an early age and it is not known whether this mutation affected dopamine neurons in the SNpc, although basal ganglia degeneration was reported.^66^ BNIP3 and NIX are constitutively degraded by the proteasome. We found several proteasomal subunits to have significantly lower gene expression in *SNCA*-*Trp* iPSC-DANs (Fig. 3), and also observed reduced proteasome activity (Fig. 2). This is in line with multiple studies reporting impaired proteasome expression and activity in the substantia nigra of PD post-mortem brains.^67–70^ Furthermore, nigral injection of a proteasome inhibitor in mice induced PD pathology including striatal dopamine decrease, α-synuclein accumulation and loss of TH^+^-neurons in the substantia nigra, while variants in the proteasome regulator PSMF1 were recently described to cause early-onset parkinsonism with mitochondrial dysfunction.^71–72^ Pharmacological or genetic proteasome inhibition increased mitophagy and impaired mitochondrial function, in line with our findings.^73–74^ Together, these results highlight the importance of the UPS in PD and connects this to the basal mitophagy axis driven by BNIP3 and NIX.

We have previously observed reduced mitochondrial protein expression in dopamine neurons in *SNCA-OVX* mice, which overexpress the human *SNCA* gene, compared to WT.^6^ In accordance, the scRNA-seq results indicate that mitochondrial genes are broadly downregulated in *SNCA-Trp* iPSC-DANs (Fig 3). This may be interpreted as a response to elevated mitophagy induced by α-synuclein to save cellular resources, although that would require further validation. Of note, mitochondrial gene expression was broadly upregulated in the midbrains of *Snca-/-*mice.^75^ Ultimately, heightened mitophagy may lead to mitochondrial depletion. Loss of mitochondrial mass has been reported in α-synuclein, Parkin, LRRK2, FBXO7 and MitoPark mouse models of PD.^14, 76–81^ In combination with the finding that dopamine neurons have lower mitochondrial mass in the vulnerable SNpc^82–85^ compared to the relatively protected neighbouring ventral tegmental area (VTA), this may provide a mechanistic rationale for the selective vulnerability of these neurons: when pressured to perform more mitophagy, SNpc neurons risk losing too many functional mitochondria. Collectively, the loss of mitochondrial mass may represent a shared driver of PD pathology and suggests that current therapeutic strategies aiming to increase mitophagy on its own may be detrimental.^86^

α-Synuclein is a well-described anionic lipid binding protein, which is mediated through its *N*-terminal domain.^33, 87^ We confirmed that recombinant α-synuclein binds specific anionic lipids including PA, CL and phosphoinositides (Fig. 4D). Using whole cell global lipidomics, we observed that total CL was increased, while total PA levels were depleted in *SNCA-Trp* iPSC-DANs. Of note, brain CL levels were reported to be decreased in *Snca-/-*mice.^88^ We found multiple lipid-related genes to be differentially expressed in *SNCA-Trp* iPSC-DANs compared to controls (Fig 3). This included decreased expression of the phospholipase PLD6, which converts CL to PA on the OMM and regulates DRP1 mitochondrial fission activity.^34, 36^ Overexpression of the α-synuclein *N*-terminal domain has been shown to be essential for eliciting DRP1-mediated mitochondrial fragmentation.^10^ Furthermore, elevated levels of DRP1 have been observed in PD patient brain and genetic inactivation of DRP1i was neuroprotective in the MPTP and *Pink-/-* mouse models.^89–90^ We and others have also observed increased p616-DRP1 expression in human *SNCA-OVX* mice.^6, 90^ These findings prompted us to investigate DRP1 blockade as a strategy to reverse the observed mitophagy and mitochondrial fragmentation phenotypes. Prior studies have opted for the DRP1 inhibitor Mdivi-1, however Mdivi-1 is a suspected antioxidant and its specificity for inhibiting DRP1 is unclear.^91–92^ Here, we showed that the DRP1 inhibitor MB-0223 reversed the increased mitophagy phenotype in *SNCA-Trp* iPSC-DANs back to control levels and improved mitochondrial morphology, intracellular ATP and calcium release (Fig. 5). Notably, we found that BNIP3 and NIX were decreased after treatment with MB-0223, which we consider to be due to the slowing of mitophagy which allows for degradation of these mitophagy receptors by the UPS. The combined results provide evidence that dampening mitophagy may provide therapeutic benefit in synuclein-driven PD.

Finally, we utilised the MEA platform to assess neuronal electrophysiological activity and overall health, which indicated that aged (>D60) *SNCA-Trp* iPSC-DANs showed impaired firing and bursting. This is in agreement with our previous work using the *SNCA-OVX* mouse model where we observed reduced neuronal firing in aged mice.^6^ A recent study in *SNCA-Trp* midbrain organoids also showed a similar loss in neuronal activity after 50 days in culture.^57^ Treatment with MB-0223 reversed the neuronal activity deficit in *SNCA-Trp* iPSC-DANs (Fig 6), confirming that mitochondrial dysfunction caused by increased α-synuclein may underlie and precede neuronal dysfunction.

Taken together, we have demonstrated that increased α-synuclein expression increases basal mitophagy in Parkinson’s iPSC-DANs through mechanisms involving increased BNIP3 and NIX and cardiolipin leading to loss of mitochondrial mass. DRP1 inhibition reversed this increase in mitophagy and rescued a range of disease-relevant cellular phenotypes highlighting a potential therapeutic route to modulate mitophagy in PD.

## Acknowledgments

E.D.M. was supported by a NWO Rubicon Fellowship (019.202EN.048). The work was supported by a National Institute for Health Research-Medical Research Council Dementias Platform UK Equipment Award (MR/M024962/1) to R.W.-M. This work was supported by the Medical Research Council grant (MR/Y014987/1) to B.J.R. C.W. was supported by the Medical Research Council, UK. A.K. and B.J. are supported by the Wellcome Trust (grant number 226800/Z/22/Z) and the NIHR Cambridge Biomedical Research Centre (NIHR203312).

## Author contributions

E.D.M., W.M., M.C.C. and M.L. differentiated iPSC lines and generated experimental data; B.V. performed bioinformatics analysis; B.J. and A.K. undertook lipidomics analysis; M.R., J.M.-S. and C.W. undertook single cell sequencing and data quality control; B.J.R. and R.W.-M. supervised the experimental work. E.D.M prepared the manuscript, B.J.R. and R.W.-M. edited the manuscript, and all authors approved the final version.

## Materials and Methods

### Chemicals

- MB0223 (Bio-Techne R&D Systems, 6799)
- MLN4924 (Pevonedistat) (Cambridge Bioscience Ltd, 15217)
- Epoxomicin (Abcam plc, ab144598)
- Carbonyl cyanide 4-(trifluoromethoxy)phenylhydrazone (CCCP) (Merck Life Science UK Ltd, C2759)
- 6-Cyano-7-nitroquinoxaline-2,3-dione (CNQX) (Tocris, 0190)
- D-2-Amino-5-phosphonopentanoic acid (D-AP5) (Abcam plc, ab120003)
- NeuroFluor NeuO (Stemcell Technologies UK Ltd, #01801)
- Mtphagy mitophagy dye (Dojindo, MD01)
- Lysotracker Deep Red (Invitrogen, L12492)
- BiotrackerATP-Red (Merck Life Science UK Ltd, SCT045)
- Fura2-AM (Molecular Devices, MLDVR8198 Fura-2 QBT™ Calcium Bulk Kit)
- NucBlue™ Live ReadyProbes™ Reagent (Invitrogen, R37605)
- DAPI (Merck Life Science UK Ltd, MBD0015)
- Suc-LLVY-AMC (Bio-Techne R&D Systems, S-280-05M)
- Doxycycline hyclate (Merck Life Science UK Ltd, D9891)
- Glutamine (Life Technologies, 25030024)
- Penicillin/streptomycin (Life technologies, 15140122)
- Antibiotic-antimycotic (Life Technologies Ltd, 15240096)
- Laminin from Engelbreth-Holm-Swarm murine sarcoma basement membrane (mouse, Merck Life Science UK Ltd, L2020)
- Biolaminin 521 LN (Biolamina AB, LN521-05)
- Poly-L-ornithine (Merck Life Science UK Limited, P4957)
- Recombinant human/murine/rat brain-derived neurotrophic factor (BDNF) (Peprotech, 450-02)
- Recombinant human glial-derived neurotrophic factor (GDNF) (Peprotech, 450-10)
- Recombinant human transforming growth factor-beta 3 (TGFβ3) (Peprotech, 100-36E)
- N6,2’-O-Dibutyryladenosine-3’,5’-cyclic monophosphate sodium salt (dbcAMP, Biosynth, ND07996)
- CHIR99021 (Bio-Techne R&D Systems, 4423)
- DAPT γ-secretase inhibitor (Abcam plc, ab120633)
- SB 431542 (Apex Bio, A8249)
- Y27632 dihydrochloride ROCK inhibitor (Apex Bio, A3008)
- Recombinant human fibroblast growth factor 8a (FGF8a) (Cambridge Bioscience, Z03172-1)
- Geltrex (Life Technologies, A1413302)
- hESC-qualified Matrigel (SLS, 354277)
- Accutase (Life Technologies, A1110501)
- MEM NEAA (Life Technologies, 11140035)
- β-Mercaptoethanol (Gibco, 31350010)
- Purmorphamine (Bio-Techne R&D Systems, 4551)
- Recombinant human Sonic Hedgehog (SHH) (Bio-Techne R&D Systems, 1845-SH-500)
- Ascorbic acid (Merck Life Science UK Ltd, A4544)
- LDN193189 hydrochloride (Merck Life Science UK Ltd, SML0559)
- Mitomycin C (Abcam plc, ab120797)
- Normal donkey serum (Jackson Immuno, 017-000-121)
- CellTiter-Glo 2.0 cell viability assay (Promega, G9242)
- Pierce bicinchoninic acid (BCA) protein assay kit (Thermo Scientific, 23225)
- Amersham ECL Prime Western blotting detection reagent (VWR International, RPN2236)
- Recombinant full length human alpha-synuclein (20 nM, Stratech, S-1001-2-RPE)
- Bovine serum albumin (Merck Life Science UK, A9418)
- Dimethyl sulfoxide (DMSO) (Merck Life Science UK, D2438)

### Media and cells

- mTeSR1 (Stemcell Technologies, 85850)
- mTeSR Plus (Stemcell Technologies, 100-0276)
- Neurobasal (Life Technologies Ltd, 21103049)
- Neurobasal w/o phenol red (Life Technologies Ltd, 12348017)
- BrainPhys (Stemcell Technologies UK Ltd, 05790)
- KO DMEM (Life Technologies Ltd, 10829018)
- KnockOut™ Serum Replacement (Life Technologies, 10828010)
- B27 supplement (Life Technologies Ltd, 17504044)
- N2 supplement (Life Technologies Ltd, 17502048)
- Accutase (Stemcell Technologies UK Ltd, 07920)
- MEM Non-essential amino acids solution (100x) (NEEA) (Gibco, 11140050)
- Rat primary cortical astrocytes (Gibco, N7745100)

### Primary antibodies

- Anti-microtubule associated protein 2 (MAP2) chicken polyclonal (ICC 1:1000; Abcam, ab92434)
- Anti-microtubule associated protein 2 (MAP2) chicken polyclonal (ICC 1:1000, Merck Milliore, AB5543)
- Anti-tyrosine hydroxylase (TH) rabbit polyclonal (ICC 1:500; Millipore, AB152)
- Anti-tyrosine hydroxylase (TH) sheep polyclonal (ICC 1:500; Millipore, AB1542)
- Anti-alpha-synuclein rabbit monoclonal [MJFR1] (ICC 1:1000, Abcam, ab138501)
- Anti-pS129-alpha-synuclein [EP1536Y] (ICC: 1:500, Abcam, ab51253)
- Anti-TOM20 mouse monoclonal [F10] (ICC 1:500; Santa Cruz Biotechnology, sc-17764)
- Anti-BNIP3 mouse monoclonal [ANa40] (ICC 1:500, Novus Biologics, NB100-1543)
- Anti-NIX rabbit polyclonal (ICC 1:500, Abcam, ab8399)
- Anti-pS65Ub rabbit monoclonal [E2J6T] (ICC 1:500, Cell Signalling, #62802)
- Anti-pS616-DRP1 rabbit monoclonal [D9A1] (ICC 1:500, Cell Signaling, #4494)
- Anti-VGLUT2 rabbit polyclonal (ICC 1:500, Synaptic Systems, 135 403)

### Secondary antibodies

- Donkey anti-mouse Alexa Fluor 647 (ICC 1:500, Invitrogen, A31571)
- Donkey anti-mouse Alexa Fluor 488 (ICC 1:500, Invitrogen, A21202)
- Donkey anti-rabbit Alexa Fluor 647 (ICC 1:500, Invitrogen, A31573)
- Donkey anti-rabbit Alexa Fluor 555 (ICC 1:500, Invitrogen, A31572)
- Donkey anti-rabbit Alexa Fluor 488 (ICC 1:500, Invitrogen, A21206)
- Donkey anti-chicken Alexa Fluor 488 (ICC 1:500, Invitrogen, A21206)
- Donkey anti-sheep Alexa Fluor 647 (ICC 1:500, Invitrogen, A21448)
- Goat-anti-rabbit-HRP (1:5000, Biorad, 1706515)

### Culture of iPSC-DANs and differentiation into dopaminergic neurons

The generation and characterization of iPSCs from healthy individuals or PD patients carrying the *SNCA-Trp* mutation were performed as previously described.^5^ iPSC lines were routinely cultured on Matrigel (BD Biosciences) in mTeSR1 or mTeSR+ (StemCell Technologies) and expanded by colony passaging. Differentiation of dopaminergic neurons was based on a previously described modified Kriks’ protocol.^11, 93^

Briefly, two days prior to neuronal induction, iPSCs were dissociated to single cells using Accutase and plated at a density of 1.5 x 10^5^ cells/cm^2^ on Geltrex-coated wells (Life Technologies, A1413302). iPSCs were patterned for 10 days to become ventral midbrain precursor cells. After patterning (DIV 10), cells were expanded for 19 days in KSR and NNB medium (1:3) supplemented with LDN193189 (100 nM, Merck) and CHIR99021 (3 μM, Tocris). Cells were passaged four times 2:3 or 1:2 with Accutase depending on cell density on days 10 + 1, 10 + 7, 10 + 13 and 10 + 19 and plated with 10 μM Y-27632 onto Geltrex coated plates. Cells were frozen at DIV 10 + 13 or D10 + 19 in 1:3 of KSR:NNB medium supplemented with 10 % (v/v) DMSO. For KSR medium: From a total of 500 mL KO DMEM, 90 mL was removed and 75 mL KSR serum replacement, 5 mL β-mercaptoethanol (50 mM), 5 mL MEM non-essential amino acids solution, and 5 mL L-glutamine (200 mM) were added. NNB medium: 500 mL Neurobasal with 5 mL B27 supplement, 2.5 mL N2 supplement, and 5 mL L-glutamine (200 mM).

Following the expansion of ventral midbrain precursor cells (DIV 11), media was changed to NB medium containing Neurobasal medium, 10 mL B27 and 2 mM L-glutamine and differentiated into dopaminergic neuronal cultures. Cells were replated at DIV 20 followed by 15 days of maturation. All experiments were carried out with 3-4 iPSC lines per genotype between DIV 35-50 and data was averaged from 2-4 independent differentiations unless otherwise indicated.

### mt-Keima assay

The mt-Keima assay was performed as previously described.^11^ iPSC-DANs (30K/well) plated on D20 on PLO/Biolamina-coated 96W half-area plates were transduced with doxycyclin-inducible mt-mKeima lentivirus (pLEX-TRE3G-mt-mKeima-pPGK-rtTA3 vector, MOI of 2) and matured to D35-D50. 48 h before imaging, doxycycline (1 µg/mL, Merck Life Science UK Ltd, D9891) and NucBlue™ Live ReadyProbes™ Reagent (Invitrogen, R37605) were added in NB maturation media without phenol red. Treatments were included together with doxycycline for 48 h (MB-0223 (1, 3, 10 µM), MLN4924 (1 µM), epoxomicin (5 nM) or DMSO 0.1%). CCCP (5 µM) was added 4 h before imaging. Live images were taken on an Opera Phenix™ High Content Screening System (Perkin Elmer) using a 40x objective with mKeima excitation at 425 nm (green) and 561 nm (red) and emission at 570-630 nm, and the DAPI channel for identification of nuclei. Images were analysed using Harmony^TM^ software (Perkin Elmer) to provide ratio images of red and green mt-mKeima signal and mitochondrial morphology (green mitochondria only). The mitophagy index is defined as the ratio of red/green total mt-mKeima area. Red only spots (mitolysosomes) were identified by substracting the green from the red channel.

### Immunocytochemistry

iPSC-DANs (30K/well) plated on D20 on PLO/Biolamina-coated 96W half-area plates were matured to D35-D50. Media was removed and pre-warmed 4% paraformaldehyde in PBS was added for 10 min. The cells were washed with PBS 3x. Fixed cells were incubated with permeabilisation buffer (0.1% v/v Triton-X100 in PBS) for 6 minutes, followed by blocking with blocking solution (10% normal donkey serum, 0.1% v/v Triton-X100 in PBS). Primary antibodies were incubated in blocking solution overnight at 4 °C, followed by 3x PBS washes. Secondary antibodies were incubated together with DAPI (1 µg/mL, Merck Life Science UK Ltd, MBD0015) for 1 h at room temperature. Cells were washed with PBS 2x and imaged on an Opera Phenix™ High Content Screening System (Perkin Elmer). Primary antibodies used: anti-microtubule associated protein 2 MAP2 (1:500; Abcam, chicken polyclonal, ab92434), anti-microtubule associated protein 2 MAP2 (1:500, Merck Millipore, chicken polyclonal, AB5543), anti-tyrosine hydroxylase TH (1:500; Millipore, rabbit polyclonal, AB152), anti-tyrosine hydroxylase TH (1:500; Millipore, sheep polyclonal, AB1542), anti-alpha-synuclein (1:1000, Abcam, rabbit monoclonal MJFR1, ab138501), anti-alpha-synuclein pS129 (1:500, rabbit monoclonal EP1536Y, Abcam, ab51253), anti-TOM20 (1:500; Santa Cruz Biotechnology, mouse monoclonal F-10, sc-17764), anti-BNIP3 (1:500, Novus Biologics, mouse monoclonal ANa40, NB100-1543), anti-BNIP3L/NIX (1:500, abcam, rabbit polyclonal, ab8399), anti-ubiquitin phosphoserine-65 pS65Ub (1:500, Cell Signalling, rabbit monoclonal E2J6T, #62802). p616-DRP1 (1:500, Cell Signaling, rabbit monoclonal D9A1, #4494). VGLUT2 (1:500, Synaptic Systems, rabbit polyclonal, 135 403). Secondary antibodies used: donkey anti-mouse Alexa Fluor 647 (Invitrogen, A31571), donkey anti-mouse Alexa Fluor 488 (Invitrogen, A21202), donkey anti-rabbit Alexa Fluor 647 (Invitrogen, A31573), donkey anti-rabbit Alexa Fluor 555 (Invitrogen, A31572), donkey anti-rabbit Alexa Fluor 488 (Invitrogen, A21206), donkey anti-chicken Alexa Fluor 488 (Invitrogen, A21206), donkey anti-sheep Alexa Fluor 647 (Invitrogen, A21448).

### Mtphagy mitophagy assay

iPSC-DANs (30K/well) plated on D20 on PLO/Biolamina-coated 96W half-area plates were matured to D35-D50 for imaging. Cells were treated with Mtphagy mitophagy dye (50 nM final concentration, 48 h, Dojindo, MD01) in NB maturation media without phenol red. 1 h before imaging the media was refreshed to which was added NeuroFluor NeuO (1 µM, Stemcell Technologies UK Ltd, # 01801), Lysotracker Deep Red (50 nM, Invitrogen, L12492) and NucBlue live (Life Technologies Ltd, R37605). Optionally, bafilomycin A1 (500 nM, Merck Life Science UK Ltd, B1793) was added 3 h before imaging to assess mitophagy flux. Live images were taken on an Opera Phenix™ High Content Screening System (Perkin Elmer) using a 63x objective for Mtphagy (ex. 488 nm; em. 650-760 nm), NeuO (ex. 488 nm; em. ), Lysotracker Deep Red (ex. 647 nm; em. and 561 nm (red) and emission at 570-630 nm, and the DAPI channel for identification of nuclei. Images were analysed using Harmony^TM^ software (Perkin Elmer)

### BiotrackerATP assay

iPSC-DANs (30K/well) plated on D20 on PLO/Biolamina-coated 96W half-area plates were matured to D35-D50 for imaging. Cells were treated with DMSO or MB-0223 (3 µM, 48 h, Bio-Techne, 6799) in NB maturation media without phenol red. 1 h before imaging the media was refreshed to which was added BiotrackerATP-Red (5 µM final concentration, Merck Life Science UK Ltd, SCT045), NeuroFluor NeuO (1 µM, Stemcell Technologies UK Ltd, # 01801), NucBlue live (Life Technologies Ltd, R37605), and included MB-0223 (3 µM) or DMSO. Cells were washed with PBS (1x) and imaged in NB maturation media without phenol red which included MB-0223 (3 µM) or DMSO.

### ATP production assay

iPSC-DANs (30K/well) plated on D20 on PLO/Geltrex-coated 96W half-area plates were matured to D35-D50. Media was removed and cells were mixed with 100 µL/well CellTiter-Glo 2.0 Cell Viability Assay (Promega, G9242) diluted in PBS (1:1) and incubated for 10 min. Then, 90 µL of substrate medium were transferred to a 96-well white plate and incubated for 25 minutes at room temperature, shielded from light. To measure ATP production, luminescence was measured PHERAstar FSX plate reader (BMG Labtech) and measurements were normalised to cell number. Cell number was measured prior to the start of the assay by adding Hoechst 33342 nuclear marker (NucBlue™ Live ReadyProbes™ Reagent, R37605, Invitrogen) to cells and imaging live (Opera Phenix™ High Content Screening System, Perkin Elmer).

### Fura2-AM calcium release assay

The Fura2-AM calcium release assay was performed as previously described.^51^ iPSC-DANs (30K/well) plated on D20 on PLO/Geltrex-coated 96W half-area plates were matured to D35-D50 for imaging. Cells were treated with DMSO or MB-0223 (3 µM, 48 h, Bio-Techne, 6799) in NB maturation media. Fura-2AM (Molecular Devices, MLDVR8198 Fura-2 QBT™ Calcium Bulk Kit) was reconstituted to 2x in Ca^2+^/Mg^2+^-free HBSS supplemented with 20 mM HEPES according to the manufacturer’s instructions. This was diluted to 1x final concentration using NB maturation media with DMSO or MB-0223 (3 μM). Cells were incubated with the Fura-2AM solution for 1 h at 37 °C, 5% CO_2_ and then ratiometric signals (ex 340/380 nm, em 510 nm) were acquired on a FlexStation 3 Multi-Mode Microplate Reader (Molecular Devices) at 37 °C every 4 s for 100 s. Ionomycin (final concentration 5 μM) was injected after a baseline of 28 s.

For the analysis, the 340/380 ratio was computed, and then the baseline was subtracted from all the time points to obtain a normalized trace for each well. The maximum intensity (peak amplitude) of the normalized trace was found and plotted.

### Multi-electrode array recordings

Axion Biosystems Cytoview 48-well MEA plates (16 electrodes/well, M768-tMEA-48W) were coated two days before cell plating (D18) with poly-L-ornithine (8 µL droplet/ well, Merck Life Science UK, P4957, 0.01% solution) and incubated overnight at 37 °C. PLO was removed and Biolaminin (8 µL droplet/well, 25 µg/mL diluted in DPBS++, Biolamina, LN521) was added and again incubated overnight at 37 °C. On the day of plating (D20), 35,000 iPSC-DANs in 6 µL Neurobasal maturation media with ROCK inhibitor were spotted on the recording electrodes and allowed to settle for 1 h at 37 °C followed by careful addition of maturation media with ROCK inhibitor (200 µL/well). After 2 days (D22), cultures were treated with mitomycin C (1 µg/ml, 1 h) in Neurobasal maturation media to remove proliferative cells. On D24, media was removed and rat primary cortical astrocytes (Gibco, N7745100) were added (1.16 * 10^6^ cells/well). After the astrocytes had settled for 1 h, 200 µL/well Neurobasal maturation media was added. From D26 half media changes (100 µL per well) were carried out three times a week using BrainPhys + B27 maturation media with the addition of laminin (2 µg/mL, Merck Life Science UK, L2020). MEA recordings (2 min, 37 °C and 5% CO2, sampling frequency = 12.5 kHz) were acquired at least once a week on a Maestro PRO (Axion Biosystems) using AxIS Navigator v3.6.2.2 from D30 onwards to assess neuronal firing activity. Plates were equilibrated in the Maestro PRO for 10 min prior to recording. An adaptive threshold spike detector set to 5.5 standard deviations was applied to the raw data. RAW files were analysed with AxIS Navigator to produce metric files describing firing and bursting activity. The following parameters were used: Active electrode criterion = 5 spikes/min; Burst detection: inter-spike interval threshold, maximum inter-spike interval = 100 ms, minimum number of spikes = 5. Weighted mean firing rate was calculated by averaging the mean firing rate across only active electrodes. Further analysis of the metric files was performed with AxIS Metric Plotting Tool v2.4.4 and Excel, and testing for statistical significance with Graphpad Prism v8. 3 healthy control and 3 *SNCA-Trp* iPSC lines were used with 8 replicates wells per line and data were averaged per iPSC line over 3 independent differentiations. Treatment with MB-0223 (3 µM) or DMSO control was performed after recording a baseline followed by a half feed (100 µL/well). MB-0223 (3 µM) or DMSO control was included in half media changes every 2-3 days. For the MB-0223 treatment four replicate wells per iPSC line per condition were used.

### Lipid dot blot assay

Membrane lipid strip dot blots (Echelon Bioscience, P-6002-2PK) were blocked with 3% BSA (Merck Life Science UK, A9418) in PBS and 0.05% Tween-20 at room temperature for 1 h. The lipid strip was treated with recombinant full length human alpha-synuclein (20 nM, Stratech, S-1001-2-RPE) in PBS overnight at 4 °C. After three washes with PBS + Tween-20, primary anti-alpha-synuclein antibody (1:5000, MJFR1, Abcam, ab138501) in 3% BSA in PBS + Tween-20 was incubated at room temperature for 1 h. Following three washes with PBS + Tween-20, goat-anti-rabbit-HRP (1:5000, Biorad, 1706515) in 3% BSA in PBS + Tween-20 was incubated at room temperature for 1 h. Chemiluminescence was visualised using Amersham ECL Prime Western blotting detection reagent (VWR International, RPN2236). Data from two lipid strips was averaged and quantified in Graphpad v8.

### Lipidomics

For whole cell lipid measurements 5*10^6^ iPSC-DANs per cell line were plated on D20 in 6W multi-well plates and matured to D40. The cells were harvested in 1 mL PBS and 100 µL cell suspension was used for normalisation. The remaining cells in the 0.9 mL suspension were pelleted by centrifugation (400 g, 10 min, 4 °C), the supernatant was discarded and the cell pellets were stored at -80 °C until lipid extraction. The 100 µL cell suspension for normalisation was lysed using a probe sonicator (10 s, 10% amplitude) and protein concentrations were determined using a Pierce bicinchoninic acid (BCA) protein assay kit (Thermo Scientific, 23225).

The lipid extraction protocol was performed as described previously.^94^ Briefly, 650 µL of chloroform was added to each sample, followed by thorough mixing. The samples were then homogenised using a Bioprep 24-1004 homogeniser (Allsheng, Hangzhou City, China) run at speed: 4.5 m/s, time: 30 seconds for 2 cycles. Then 100 µL of the LIPID internal standard (1-10 µM in methanol), 100 µL of the CARNITINE internal standard (5 µM in methanol) and 150 µL of methanol was added to each sample, followed by thorough mixing. Then, 400 µL of acetone was added to each samples. The samples were vortexed and centrifuged for 10 minutes at ∼20,000 g to pellet any insoluble material. The supernatant was pipetted into separate 2 mL screw cap amber-glass auto-sampler vials (Agilent Technologies, Cheadle, United Kingdom). The organic extracts were dried down to dryness using a Concentrator Plus system (Eppendorf, Stevenage, United Kingdom) run for 60 minutes at 60 °C.

For lipidomics LC-MS analysis: the samples were reconstituted in 100 µL of 2:1:1 (propan-2-ol, acetonitrile and water, respectively) then thoroughly vortexed. The reconstituted sample was transferred into a 250 μL low-volume vial insert inside a 2 mL amber glass auto-sample vial ready for liquid chromatography with mass spectrometry detection (LC-MS) lipidomics analysis. Full chromatographic separation of intact lipids was achieved using Waters Acquity H-Class HPLC System (Waters, Hertfordshire, United Kingdom) with the injection of 10 µL onto a Waters Acquity UPLC® CSH C18 column; 1.7 µm, I.D. 2.1 mm X 50 mm, maintained at 55 degrees Celsius. Mobile phase A was 6:4, acetonitrile and water with 10 mM ammonium formate. Mobile phase B was 9:1, propan-2-ol and acetonitrile with 10 mM ammonium formate. The flow was maintained at 500 µL per minute through the following gradient: 0.00 minutes_40% mobile phase B; 1.5 minutes_40% mobile phase B; 8.00 minutes_99% mobile phase B; 10.00 minutes_99% mobile phase B; 10.10 minutes_40% mobile phase B; 12.00 minutes_40% mobile phase B. The sample injection needle was washed using 9:1, propan-2-ol and acetonitrile [strong wash] and 2:1:1 (propan-2-ol, acetonitrile and water) [weak wash]. The mass spectrometer used was the Thermo Scientific Exactive Orbitrap with a heated electrospray ionization source (Thermo Fisher Scientific, Hemel Hempstead, United Kingdom). The mass spectrometer was calibrated immediately before sample analysis using positive and negative ionisation calibration solution (recommended by Thermo Scientific). Additionally, the mass spectrometer scan rate was set at 4 Hz, giving a resolution of 25,000 (at 200 m/z) with a full-scan range of m/z 100 to 1,800 with continuous switching between positive and negative mode.

Lipid levels are expressed in nmol/mg protein after correction for protein concentration. Data represent the mean ± SEM of 3 healthy control and 3 *SNCA-Trp* iPSC-DANs lines for one differentiation. For each iPSC-DAN line 3 technical replicates were averaged.

### Proteasome activity assay

The proteasome activity assay was adapted from a previous study.^95^ iPSC-DANs (30K/well) plated on D20 on PLO/Geltrex-coated 96W half-area plates were matured to D35-D50 and treated with epoxomicin (1 µM, 1 h) or DMSO in NB maturation media. After removal of the media, cells were treated with assay buffer (20 mM Tris, 5 mM MgCl_2_, 1 mM DTT, 1 mM ATP, pH = 7.4 in MilliQ) containing 100 μM suc-LLVY-AMC (Bio-Techne R&D Systems, S-280-05M). Fluorescence (λ ex / em = 355 / 450 nm) was measured immediately on a PHERAstar FSX plate reader (BMG Labtech) every 2 min for 1 h at 37 °C. The increase in fluorescence per minute was used to calculate the slope (= proteasome activity) of the sample. Epoxomicin treated wells were used as background correction and set to 0%. Data represent the mean ± SEM of 3 healthy control and 3 *SNCA-Trp* iPSC-DANs lines for three independent differentiations.

### Single Cell RNA sequencing

#### Harvesting and scRNA-seq of iPSC-DANs

Mature iPSC-DANs were dissociated with papain and Accutase following a previously described protocol^96^, fixed in 80% MeOH^97^ and stored at -80 °C. For rehydration, fixed cells were equilibrated to 4 °C and subjected to two rounds of centrifugation at 1,000 g for 5 min at 4 °C, followed by resuspension in ice-cold PBS supplemented with 0.01% BSA, 5 mM DTT, 400 U/ml recombinant RNase inhibitor. Following rehydration, single cells were dispensed into the nanowells of an ICELL8 350v Chip using the ICELL8 cx Single-Cell System (Takara Bio) and ICELL8 cx CELLSTUDIO v2.5 Software, before selection of candidate wells with the ICELL8 cx CellSelect v2.5 Software. cDNA libraries were constructed using the SMART-Seq Pro Application kit and sequenced on an Illumina Novaseq instrument (PE 100, ∼0.5-1 million read pair / cell).

#### Preprocessing of raw sequencing data

Raw base call (BCL) files were converted to FASTQ format and demultiplexed by sample index using bcl2fastq2. Sample-specific FASTQ files were then split into cell barcode-level FASTQ files using Cogent NGS Analysis Pipeline 2.0.1 (cogent demux with: --type_of_experiment ICELL8_FLA, --undetermined_fq). Raw sequencing reads were quality checked with FastQC / MultiQC and adaptor trimming was performed with Trim Galore (--stringency 3, --trim-n).

#### Transcript quantification with Salmon

A decoy-aware transcriptome was constructed by concatenating the human transcriptome with the primary genome assembly (GENCODE release 44, GRCh38.p14) and indexed using Salmon with k-mers of length 31 (salmon index with: --gencode). Trimmed reads were then mapped to the decoy-aware transcriptome by selective alignment and quantified using Salmon (salmon quant with: --libType A, -- gcBias, --seqBias, --allowDovetail, --dumpEq, --writeUnmappedNames). Estimated gene counts were generated from transcript abundance estimates scaled using the average transcript length over samples and then the library size using the tximport R package (tximport::tximport() with countsFromAbundance = “lengthScaledTPM”). Downstream analyses were performed on cells with at least 20,000 raw counts and 3,000 detected genes, and mitochondrial reads accounting for no more than 20% of all TPMs. Genes detected with at least 5 raw counts in at least 1% of the retained cells were kept for downstream analyses.

#### Cell clustering and annotation

The 2,000 most variable genes were identified using the Scanpy Python library, providing the raw count matrix as input and line ID as the batch key (sc.pp.highly_variable_genes() with flavor = “seurat_v3”, n_top_genes = 2000, layer = <raw counts> and batch_key = <line ID>). To remove batch effects on cell-to-cell distances in a low-dimensional space, the raw count data was restricted to these highly variable genes and integrated using the scVI model from the scvi-tools Python library, with line ID as the batch covariate (scvi.model.SCVI.setup_anndata() with layer = <raw counts> and categorical_covariate_keys = <line ID>). The resulting scVI latent representation served to compute a k-nearest neighbour graph of cells (sc.pp.neighbors() with use_rep = “X_scVI”), which were then clustered using the Leiden algorithm (sc.tl.leiden() with resolution = 0.6). Clusters were annotated based on marker gene expression, which revealed three major cell types: neuron, astrocyte and fibroblast-like, characterized by high expression of SNAP25 / TH, AQP4 and COL1A1, respectively.

#### Differential gene expression analysis

Differential gene expression analysis (DGEA) was performed at the single-cell level using the MAST R package. The dataset was restricted to cells annotated as neurons and to protein-coding genes detected (TPM > 0) in at least 20% of these cells. TPM estimates were log2-transformed after addition of a pseudocount of one and used as input expression values. The effect of genotype (*SNCA-Trp* vs control, control as reference) was estimated with a hurdle model of the form ∼ CDR + genotype + (1 | line ID), where CDR denotes the cellular detection rate (the scaled number of genes detected per cell). Because the model included a random intercept for line ID, it was fitted as a generalized linear mixed-effects model (zlm() with method = “glmer”, ebayes = FALSE and strictConvergence = FALSE). Statistical significance was assessed by likelihood ratio test on the genotype coefficient. For genes whose hurdle model did not return an estimate, the log_2_ fold-change and *P*-value were recovered from a linear mixed-effects model fitted to the continuous (detection-conditional) component alone, using the same fixed and random effects. p values were adjusted across all tested genes using the Benjamini–Hochberg method, and genes with adjusted *P* < 0.05 were considered differentially expressed. Mitochondrial-related genes were identified using the MitoCarta3.0 database.^39^ Lipid-related and autophagy-related genes were identified using the annotations of the gene set enrichment analysis searching for keywords FATTY_ACID, LIPID, ADIPOGENESIS, ADIPOCYTE, ADIPOSE or AUTOPHAGY, AUTOPHAGOSOME, respectively.

#### Gene set enrichment analysis

Gene set enrichment analysis (GSEA) was conducted using the fgsea R package against gene sets from the Molecular Signatures Database (MSigDB, v2025.1.Hs): the Hallmark collection (H), the canonical pathway sub-collections KEGG_LEGACY, REACTOME and WIKIPATHWAYS (C2), and the Gene Ontology biological process sub-collection (C5, GO:BP). Genes were ranked by the product of the sign of the log_2_ fold-change estimate and the negative log_10_-transformed raw p value from the differential expression analysis [sign(log2 fold change) × −log_10_(raw p value)].

Enrichment was tested for each collection separately, considering only gene sets containing 15–500 genes (fgsea() with pathways = <collection-specific gene sets>, stats = <gene ranking metric>, minSize = 15, maxSize = 500).

## List of Supplementary Materials

Supplementary Figures 1-10

Supplementary Tables 1-3

## References

1. Malpartida, A. B.; Williamson, M.; Narendra, D. P.; Wade-Martins, R.; Ryan, B. J., Mitochondrial Dysfunction and Mitophagy in Parkinson’s Disease: From Mechanism to Therapy. Trends Biochem Sci 2021, 46 (4), 329–343.

2. Polymeropoulos, M. H.; Lavedan, C.; Leroy, E.; Ide, S. E.; Dehejia, A.; Dutra, A.; Pike, B.; Root, H.; Rubenstein, J.; Boyer, R.; Stenroos, E. S.; Chandrasekharappa, S.; Athanassiadou, A.; Papapetropoulos, T.; Johnson, W. G.; Lazzarini, A. M.; Duvoisin, R. C.; DiIorio, G.; Golbe, L. I.; Nussbaum, R. L., Mutation in the alpha-synuclein gene identified in families with Parkinson’s disease. Science 1997, 276 (5321), 2045–2047.

3. Nussbaum, R. L., Genetics of Synucleinopathies. Cold Spring Harbor Perspectives in Medicine 2018, 8 (6).

4. Singleton, A. B.; Farrer, M.; Johnson, J.; Singleton, A.; Hague, S.; Kachergus, J.; Hulihan, M.; Peuralinna, T.; Dutra, A.; Nussbaum, R.; Lincoln, S.; Crawley, A.; Hanson, M.; Maraganore, D.; Adler, C.; Cookson, M. R.; Muenter, M.; Baptista, M.; Miller, D.; Blancato, J.; Hardy, J.; Gwinn-Hardy, K., α-synuclein locus triplication causes Parkinson’s disease. Science 2003, 302 (5646), 841–841.

5. Zambon, F.; Cherubini, M.; Fernandes, H. J. R.; Lang, C.; Ryan, B. J.; Volpato, V.; Bengoa-Vergniory, N.; Vingill, S.; Attar, M.; Booth, H. D. E.; Haenseler, W.; Vowles, J.; Bowden, R.; Webber, C.; Cowley, S. A.; Wade-Martins, R., Cellular α-synuclein pathology is associated with bioenergetic dysfunction in Parkinson’s iPSC-derived dopamine neurons. Hum Mol Genet 2019, 28 (12), 2001–2013.

6. Ryan, B. J.; Bengoa-Vergniory, N.; Williamson, M.; Kirkiz, E.; Roberts, R.; Corda, G.; Sloan, M.; Saqlain, S.; Cherubini, M.; Poppinga, J.; Bogtofte, H.; Cioroch, M.; Hester, S.; Wade-Martins, R., REST Protects Dopaminergic Neurons from Mitochondrial and α-Synuclein Oligomer Pathology in an Alpha Synuclein Overexpressing BAC-Transgenic Mouse Model. J Neurosci 2021, 41 (16), 3731–3746.

7. Virdi, G. S.; Choi, M. L.; Evans, J. R.; Yao, Z.; Athauda, D.; Strohbuecker, S.; Nirujogi, R. S.; Wernick, A. I.; Pelegrina-Hidalgo, N.; Leighton, C.; Saleeb, R. S.; Kopach, O.; Alrashidi, H.; Melandri, D.; Perez-Lloret, J.; Angelova, P. R.; Sylantyev, S.; Eaton, S.; Heales, S.; Rusakov, D. A.; Alessi, D. R.; Kunath, T.; Horrocks, M. H.; Abramov, A. Y.; Patani, R.; Gandhi, S., Protein aggregation and calcium dysregulation are hallmarks of familial Parkinson’s disease in midbrain dopaminergic neurons. NPJ Parkinsons Dis 2022, 8 (1), 162.

8. Thorne, N. J.; Tumbarello, D. A., The relationship of alpha-synuclein to mitochondrial dynamics and quality control. Frontiers in Molecular Neuroscience 2022, 15.

9. Nakamura, K.; Nemani, V. M.; Azarbal, F.; Skibinski, G.; Levy, J. M.; Egami, K.; Munishkina, L.; Zhang, J.; Gardner, B.; Wakabayashi, J.; Sesaki, H.; Cheng, Y. F.; Finkbeiner, S.; Nussbaum, R. L.; Masliah, E.; Edwards, R. H., Direct Membrane Association Drives Mitochondrial Fission by the Parkinson Disease-associated Protein α-Synuclein. Journal of Biological Chemistry 2011, 286 (23), 20710–20726.

10. Krzystek, T. J.; Banerjee, R.; Thurston, L.; Huang, J.; Swinter, K.; Rahman, S. N.; Falzone, T. L.; Gunawardena, S., Differential mitochondrial roles for α-synuclein in DRP1-dependent fission and PINK1/Parkin-mediated oxidation. Cell Death Dis 2021, 12 (9), 796.

11. Williamson, M. G.; Madureira, M.; McGuinness, W.; Heon-Roberts, R.; Mock, E. D.; Naidoo, K.; Cramb, K. M. L.; Caiazza, M. C.; Malpartida, A. B.; Lavelle, M.; Savory, K.; Humble, S. W.; Patterson, R.; Davis, J. B.; Connor-Robson, N.; Ryan, B. J.; Wade-Martins, R., Mitochondrial dysfunction and mitophagy defects in LRRK2-R1441C Parkinson’s disease models. Hum Mol Genet 2023, 32 (18), 2808–2821.

12. Basak, B.; Holzbaur, E. L. F., Mitophagy in Neurons: Mechanisms Regulating Mitochondrial Turnover and Neuronal Homeostasis. Journal of Molecular Biology 2025, 437 (18), 169161.

13. Chinta, S. J.; Mallajosyula, J. K.; Rane, A.; Andersen, J. K., Mitochondrial alpha-synuclein accumulation impairs complex I function in dopaminergic neurons and results in increased mitophagy in vivo. Neuroscience Letters 2010, 486 (3), 235–239.

14. Choubey, V.; Safiulina, D.; Vaarmann, A.; Cagalinec, M.; Wareski, P.; Kuum, M.; Zharkovsky, A.; Kaasik, A., Mutant A53T α-Synuclein Induces Neuronal Death by Increasing Mitochondrial Autophagy. Journal of Biological Chemistry 2011, 286 (12), 10814–10824.

15. Ryan, T.; Bamm, V. V.; Stykel, M. G.; Coackley, C. L.; Humphries, K. M.; Jamieson-Williams, R.; Ambasudhan, R.; Mosser, D. D.; Lipton, S. A.; Harauz, G.; Ryan, S. D., Cardiolipin exposure on the outer mitochondrial membrane modulates α-synuclein. Nature Communications 2018, 9 (1), 817.

16. Lurette, O.; Martín-Jiménez, R.; Khan, M.; Sheta, R.; Jean, S.; Schofield, M.; Teixeira, M.; Rodriguez-Aller, R.; Perron, I.; Oueslati, A.; Hebert-Chatelain, E., Aggregation of alpha-synuclein disrupts mitochondrial metabolism and induce mitophagy via cardiolipin externalization. Cell Death Dis 2023, 14 (11), 729.

17. Iannielli, A.; Luoni, M.; Giannelli, S. G.; Ferese, R.; Ordazzo, G.; Fossati, M.; Raimondi, A.; Opazo, F.; Corti, O.; Prehn, J. H. M.; Gambardella, S.; Melki, R.; Broccoli, V., Modeling native and seeded Synuclein aggregation and related cellular dysfunctions in dopaminergic neurons derived by a new set of isogenic iPSC lines with SNCA multiplications. Cell Death & Disease 2022, 13 (10), 881.

18. Hui, S.; George, J.; Kapadia, M.; Chau, H.; Bariring, Z.; Earnshaw, R.; Shafiq, K.; Kalia, L. V.; Kalia, S. K., Mitophagy Upregulation Occurs Early in the Neurodegenerative Process Mediated by α-Synuclein. Molecular Neurobiology 2024, 61 (11), 9032–9042.

19. Riederer, P.; Berg, D.; Casadei, N.; Cheng, F.; Classen, J.; Dresel, C.; Jost, W.; Krüger, R.; Müller, T.; Reichmann, H.; Rieß, O.; Storch, A.; Strobel, S.; van Eimeren, T.; Völker, H.-U.; Winkler, J.; Winklhofer, K. F.; Wüllner, U.; Zunke, F.; Monoranu, C.-M., α-Synuclein in Parkinson’s disease: causal or bystander? Journal of Neural Transmission 2019, 126 (7), 815–840.

20. Chu, Y.; Kordower, J. H., Age-associated increases of α-synuclein in monkeys and humans are associated with nigrostriatal dopamine depletion: Is this the target for Parkinson’s disease? Neurobiology of Disease 2007, 25 (1), 134–149.

21. Xuan, Q.; Xu, S.-L.; Lu, D.-H.; Yu, S.; Zhou, M.; Uéda, K.; Cui, Y.-Q.; Zhang, B.-Y.; Chan, P., Increase expression of α-synuclein in aged human brain associated with neuromelanin accumulation. Journal of Neural Transmission 2011, 118 (11), 1575–1583.

22. Reeve, A.; Simcox, E.; Turnbull, D., Ageing and Parkinson’s disease: Why is advancing age the biggest risk factor? Ageing Research Reviews 2014, 14, 19–30.

23. Mehellou, Y., Parkinson’s Disease: Are PINK1 Activators Inching Closer to the Clinic? ACS Medicinal Chemistry Letters 2023, 14 (7), 870–874.

24. Cao, Y.; Zheng, J.; Wan, H. Y.; Sun, Y. Q.; Fu, S.; Liu, S. S.; He, B. Y.; Cai, G. H.; Cao, Y.; Huang, H. W.; Li, Q.; Ma, Y.; Chen, S.; Wang, F. C.; Jiang, H., A mitochondrial SCF-FBXL4 ubiquitin E3 ligase complex degrades BNIP3 and NIX to restrain mitophagy and prevent mitochondrial disease. EMBO Journal 2023, 42 (13).

25. Elcocks, H.; Brazel, A. J.; McCarron, K. R.; Kaulich, M.; Husnjak, K.; Mortiboys, H.; Clague, M. J.; Urbe, S., FBXL4 ubiquitin ligase deficiency promotes mitophagy by elevating NIX levels. EMBO Journal 2023, 42 (13).

26. Nguyen-Dien, G. T.; Kozul, K. L.; Cui, Y.; Townsend, B.; Kulkarni, P. G.; Ooi, S. S.; Marzio, A.; Carrodus, N.; Zuryn, S.; Pagano, M.; Parton, R. G.; Lazarou, M.; Millard, S. S.; Taylor, R. W.; Collins, B. M.; Jones, M. J. K.; Pagan, J. K., FBXL4 suppresses mitophagy by restricting the accumulation of NIX and BNIP3 mitophagy receptors. EMBO Journal 2023, 42 (13).

27. Gao, K.; Chen, Y.; Mo, R.; Wang, C., Excessive BNIP3-and BNIP3L-dependent mitophagy underlies the pathogenesis of FBXL4-mutated mitochondrial DNA depletion syndrome. Autophagy 2024, 20 (2), 460–462.

28. Wilhelm, L. P.; Ganley, I. G., Eating your mitochondria-when too much of a good thing turns bad. EMBO Journal 2023, 42 (13), e114542.

29. Longo, A. M.; Bishnu, A.; Risiglione, P.; Montava-Garriga, L.; Cuenco, J.; Sakamoto, K.; Mackintosh, C.; Ganley, I. G., Opposing roles for AMPK in regulating distinct mitophagy pathways. Molecular Cell 2024, 84 (22).

30. Fanning, S.; Selkoe, D.; Dettmer, U., Parkinson’s disease: proteinopathy or lipidopathy? Npj Parkinsons Disease 2020, 6 (1).

31. Flores-Leon, M.; Outeiro, T. F., More than meets the eye in Parkinson’s disease and other synucleinopathies: from proteinopathy to lipidopathy. Acta Neuropathol 2023, 146 (3), 369–385.

32. Yamada, H.; Hoshino, F.; Lu, Q.; Sakane, F., Cellular phosphatidic acid sensor, α-synuclein N-terminal domain, detects endogenous phosphatidic acid in macrophagic phagosomes and neuronal growth cones. Biochem Biophys Rep 2020, 22, 100769.

33. Yamada, H.; Mizuno, S.; Honda, S.; Takahashi, D.; Sakane, F., Characterization of α-synuclein N-terminal domain as a novel cellular phosphatidic acid sensor. Febs j 2020, 287 (11), 2212–2234.

34. Adachi, Y.; Itoh, K.; Yamada, T.; Cerveny, K. L.; Suzuki, T. L.; Macdonald, P.; Frohman, M. A.; Ramachandran, R.; Iijima, M.; Sesaki, H., Coincident Phosphatidic Acid Interaction Restrains Drp1 in Mitochondrial Division. Mol Cell 2016, 63 (6), 1034–43.

35. Kameoka, S.; Adachi, Y.; Okamoto, K.; Iijima, M.; Sesaki, H., Phosphatidic Acid and Cardiolipin Coordinate Mitochondrial Dynamics. Trends Cell Biol 2018, 28 (1), 67–76.

36. Choi, S. Y.; Huang, P.; Jenkins, G. M.; Chan, D. C.; Schiller, J.; Frohman, M. A., A common lipid links Mfn-mediated mitochondrial fusion and SNARE-regulated exocytosis. Nat Cell Biol 2006, 8 (11), 1255–62.

37. Iwashita, H.; Torii, S.; Nagahora, N.; Ishiyama, M.; Shioji, K.; Sasamoto, K.; Shimizu, S.; Okuma, K., Live Cell Imaging of Mitochondrial Autophagy with a Novel Fluorescent Small Molecule. ACS Chem Biol 2017, 12 (10), 2546–2551.

38. Antico, O.; Thompson, P. W.; Hertz, N. T.; Muqit, M. M. K.; Parton, L. E., Targeting mitophagy in neurodegenerative diseases. Nature Reviews Drugs Discovery 2025, 24 (4), 276–299.

39. Rath, S.; Sharma, R.; Gupta, R.; Ast, T.; Chan, C.; Durham, T. J.; Goodman, R. P.; Grabarek, Z.; Haas, M. E.; Hung, W. H. W.; Joshi, P. R.; Jourdain, A. A.; Kim, S. H.; Kotrys, A. V.; Lam, S. S.; McCoy, J. G.; Meisel, J. D.; Miranda, M.; Panda, A.; Patgiri, A.; Rogers, R.; Sadre, S.; Shah, H.; Skinner, O. S.; To, T. L.; Walker, M. A.; Wang, H.; Ward, P. S.; Wengrod, J.; Yuan, C. C.; Calvo, S. E.; Mootha, V. K., MitoCarta3.0: an updated mitochondrial proteome now with sub-organelle localization and pathway annotations. Nucleic Acids Res 2021, 49 (D1), D1541–d1547.

40. McGuinness, W.; Ryan, B.; Guardado, C. L.; Carling, P.; Vallin, B.; Caiazza, M. C.; Heon-Roberts, R.; Kilfeather, P.; Wantling, J.; Hoffmann, J.; Aird, D.; Lofstromm, K.; Zhang, M.; Cowley, S. A.; Hirst, W. D.; Wade-Martins, R., TFEB and TFE3 have cell-type specific expression in the brain and divergent roles in neurons. bioRxiv 2025, 2025.04.22.649949.

41. Schepers, J.; Löser, T.; Behl, C., Lipids and α-Synuclein: adding further variables to the equation. Frontiers in Molecular Biosciences 2024, *Volume* 11 *-*2024.

42. Doll, S.; Proneth, B.; Tyurina, Y. Y.; Panzilius, E.; Kobayashi, S.; Ingold, I.; Irmler, M.; Beckers, J.; Aichler, M.; Walch, A.; Prokisch, H.; Trümbach, D.; Mao, G.; Qu, F.; Bayir, H.; Füllekrug, J.; Scheel, C. H.; Wurst, W.; Schick, J. A.; Kagan, V. E.; Angeli, J. P. F.; Conrad, M., ACSL4 dictates ferroptosis sensitivity by shaping cellular lipid composition. Nature Chemical Biology 2017, 13 (1), 91–98.

43. Mahoney-Sanchez, L.; Bouchaoui, H.; Boussaad, I.; Jonneaux, A.; Timmerman, K.; Berdeaux, O.; Ayton, S.; Krüger, R.; Duce, J. A.; Devos, D.; Devedjian, J. C., Alpha synuclein determines ferroptosis sensitivity in dopaminergic neurons via modulation of ether-phospholipid membrane composition. Cell Reports 2022, 40 (8).

44. Chu, C. T.; Ji, J.; Dagda, R. K.; Jiang, J. F.; Tyurina, Y. Y.; Kapralov, A. A.; Tyurin, V. A.; Yanamala, N.; Shrivastava, I. H.; Mohammadyani, D.; Wang, K. Z. Q.; Zhu, J.; Klein-Seetharaman, J.; Balasubramanian, K.; Amoscato, A. A.; Borisenko, G.; Huang, Z.; Gusdon, A. M.; Cheikhi, A.; Steer, E. K.; Wang, R.; Baty, C.; Watkins, S.; Bahar, I.; Bayir, H.; Kagan, V. E., Cardiolipin externalization to the outer mitochondrial membrane acts as an elimination signal for mitophagy in neuronal cells. Nat Cell Biol 2013, 15 (10), 1197–1205.

45. Gilmozzi, V.; Gentile, G.; Castelo Rueda, M. P.; Hicks, A. A.; Pramstaller, P. P.; Zanon, A.; Lévesque, M.; Pichler, I., Interaction of Alpha-Synuclein With Lipids: Mitochondrial Cardiolipin as a Critical Player in the Pathogenesis of Parkinson’s Disease. Front Neurosci 2020, 14, 578993.

46. Chen, Y.; Culetto, E.; Legouis, R., The strange case of Drp1 in autophagy: Jekyll and Hyde? BioEssays 2022, 44 (4), 2100271.

47. Kamerkar, S. C.; Liu, A.; Higgs, H. N., Mitochondrial fission – changing perspectives for future progress. Journal of Cell Science 2025, 138 (9).

48. Mallat, A.; Uchiyama, L. F.; Lewis, S. C.; Fredenburg, R. A.; Terada, Y.; Ji, N.; Nunnari, J.; Tseng, C. C., Discovery and characterization of selective small molecule inhibitors of the mammalian mitochondrial division dynamin, DRP1. Biochem Biophys Res Commun 2018, 499 (3), 556–562.

49. Wang, L.; Yuan, L.; Zeng, X.; Peng, J.; Ni, Y.; Er, J. C.; Xu, W.; Agrawalla, B. K.; Su, D.; Kim, B.; Chang, Y. T., A Multisite-Binding Switchable Fluorescent Probe for Monitoring Mitochondrial ATP Level Fluctuation in Live Cells. Angew Chem Int Ed Engl 2016, 55 (5), 1773–6.

50. Devine, M. J.; Kittler, J. T., Mitochondria at the neuronal presynapse in health and disease. Nature Reviews Neuroscience 2018, 19 (2), 63–80.

51. Kilfeather, P.; Khoo, J. H.; Wagner, K.; Liang, H.; Caiazza, M. C.; An, Y.; Zhang, X.; Chen, X.; Connor-Robson, N.; Shang, Z.; Wade-Martins, R., Single-cell spatial transcriptomic and translatomic profiling of dopaminergic neurons in health, aging, and disease. Cell Rep 2024, 43 (3), 113784.

52. Pré, D.; Wooten, A. T.; Biesmans, S.; Hinckley, S.; Zhou, H.; Sherman, S. P.; Kakad, P.; Gearhart, J.; Bang, A. G., Development of a platform to investigate long-term potentiation in human iPSC-derived neuronal networks. Stem Cell Reports 2022, 17 (9), 2141–2155.

53. Cramb, K. M. L.; Noor, H.; Thomas-Wright, I.; Caiazza, M. C.; Szunyogh, S.; Milosevic, I.; Beccano-Kelly, D.; Cragg, S. J.; Wade-Martins, R., Dopamine release from Parkinson’s patient-derived neurons is disrupted due to impaired synaptic vesicle loading. bioRxiv 2026, 2026.03.11.711038.

54. Kinnart, I.; Manders, L.; Heyninck, T.; Imberechts, D.; Praschberger, R.; Schoovaerts, N.; Verfaillie, C.; Verstreken, P.; Vandenberghe, W., Elevated α-synuclein levels inhibit mitophagic flux. npj Parkinson’s Disease 2024, 10 (1), 80.

55. Mazzulli, J. R.; Zunke, F.; Isacson, O.; Studer, L.; Krainc, D., α-Synuclein-induced lysosomal dysfunction occurs through disruptions in protein trafficking in human midbrain synucleinopathy models. Proc Natl Acad Sci U S A 2016, 113 (7), 1931–6.

56. Laperle, A. H.; Sances, S.; Yucer, N.; Dardov, V. J.; Garcia, V. J.; Ho, R.; Fulton, A. N.; Jones, M. R.; Roxas, K. M.; Avalos, P.; West, D.; Banuelos, M. G.; Shu, Z.; Murali, R.; Maidment, N. T.; Van Eyk, J. E.; Tagliati, M.; Svendsen, C. N., iPSC modeling of young-onset Parkinson’s disease reveals a molecular signature of disease and novel therapeutic candidates. Nature Medicine 2020, 26 (2), 289–299.

57. Serra-Almeida, C.; Jarazo, J.; Gomez-Giro, G.; Rosety, I.; Zagare, A.; Ferrante, D.; Saraiva, C.; Frangenberg, D.; Modamio-Chamarro, J.; Zuccoli, E.; Cristóvão, A. C.; Bernardino, L.; Schwamborn, J. C., Autophagy dysfunction in iPSCs-derived neurons and midbrain organoids carrying a SNCA triplication. npj Parkinson’s Disease 2026.

58. Shaltouki, A.; Hsieh, C.-H.; Kim, M. J.; Wang, X., Alpha-synuclein delays mitophagy and targeting Miro rescues neuron loss in Parkinson’s models. Acta Neuropathologica 2018, 136 (4), 607–620.

59. Shahmoradian, S. H.; Lewis, A. J.; Genoud, C.; Hench, J.; Moors, T. E.; Navarro, P. P.; Castaño-Díez, D.; Schweighauser, G.; Graff-Meyer, A.; Godie, K. N.; Sütterlin, R.; Huisman, E.; Ingrassia, A.; de Gier, Y.; Rozemuller, A. J. M.; Wang, J.; De Paepe, A.; Erny, J.; Staempfli, A.; Hoernschemeyer, J.; Grosserüschkamp, F.; Niedieker, D.; El-Mashtoly, S. F.; Quadri, M.; Van Ijcken, W. F. J.; Bonifati, V.; Gerwert, K.; Bohrmann, B.; Frank, S.; Britschgi, M.; Stahlberg, H.; Van de Berg, W. D. J.; Lauer, M. E., Lewy pathology in Parkinson’s disease consists of crowded organelles and lipid membranes. Nature Neuroscience 2019, 22 (7), 1099-+.

60. Moors, T. E.; Maat, C. A.; Niedieker, D.; Mona, D.; Petersen, D.; Timmermans-Huisman, E.; Kole, J.; El-Mashtoly, S. F.; Spycher, L.; Zago, W.; Barbour, R.; Mundigl, O.; Kaluza, K.; Huber, S.; Hug, M. N.; Kremer, T.; Ritter, M.; Dziadek, S.; Geurts, J. J. G.; Gerwert, K.; Britschgi, M.; van de Berg, W. D. J., The subcellular arrangement of alpha-synuclein proteoforms in the Parkinson’s disease brain as revealed by multicolor STED microscopy. Acta Neuropathologica 2021, 142 (3), 423–448.

61. Mahul-Mellier, A.-L.; Burtscher, J.; Maharjan, N.; Weerens, L.; Croisier, M.; Kuttler, F.; Leleu, M.; Knott, G. W.; Lashuel, H. A., The process of Lewy body formation, rather than simply α-synuclein fibrillization, is one of the major drivers of neurodegeneration. Proceedings of the National Academy of Sciences 2020, 117 (9), 4971–4982.

62. Rappe, A.; Vihinen, H. A.; Suomi, F.; Hassinen, A. J.; Ehsan, H.; Jokitalo, E. S.; McWilliams, T. G., Longitudinal autophagy profiling of the mammalian brain reveals sustained mitophagy throughout healthy aging. The EMBO Journal 2024, 43 (23), 6199–6231-6231.

63. Jiménez-Loygorri, J. I.; Villarejo-Zori, B.; Viedma-Poyatos, Á.; Zapata-Muñoz, J.; Benítez-Fernández, R.; Frutos-Lisón, M. D.; Tomás-Barberán, F. A.; Espín, J. C.; Area-Gómez, E.; Gomez-Duran, A.; Boya, P., Mitophagy curtails cytosolic mtDNA-dependent activation of cGAS/STING inflammation during aging. Nature Communications 2024, 15 (1), 830.

64. Chen, Y.; Jiao, D.; Liu, Y.; Xu, X.; Wang, Y.; Luo, X.; Saiyin, H.; Li, Y.; Gao, K.; Chen, Y.; Zhao, S. M.; Ma, L.; Wang, C., FBXL4 mutations cause excessive mitophagy via BNIP3/BNIP3L accumulation leading to mitochondrial DNA depletion syndrome. Cell Death Differ 2023, 30 (10), 2351–2363.

65. Gao, K.; Chen, Y.; Mo, R.; Wang, C., Excessive BNIP3-and BNIP3L-dependent mitophagy underlies the pathogenesis of FBXL4-mutated mitochondrial DNA depletion syndrome. Autophagy 2023.

66. Akiba, T.; Shimada, S.; Matsuda, S.; Ishida, S.; Baba, Y.; Yamashita, A.; Shoji, H.; Okazaki, Y.; Murayama, K., Siblings of FBXL4-related mitochondrial DNA depletion syndrome, leading to fatal fulminant pneumonia. Molecular Genetics and Metabolism Reports 2025, 45, 101266.

67. McNaught, K. S. P.; Belizaire, R.; Jenner, P.; Olanow, C. W.; Isacson, O., Selective loss of 20S proteasome α-subunits in the substantia nigra pars compacta in Parkinson’s disease. Neuroscience Letters 2002, 326 (3), 155–158.

68. Tofaris, G. K.; Razzaq, A.; Ghetti, B.; Lilley, K. S.; Spillantini, M. G., Ubiquitination of α-Synuclein in Lewy Bodies Is a Pathological Event Not Associated with Impairment of Proteasome Function*. Journal of Biological Chemistry 2003, 278 (45), 44405–44411.

69. Bedford, L.; Hay, D.; Devoy, A.; Paine, S.; Powe, D. G.; Seth, R.; Gray, T.; Topham, I.; Fone, K.; Rezvani, N.; Mee, M.; Soane, T.; Layfield, R.; Sheppard, P. W.; Ebendal, T.; Usoskin, D.; Lowe, J.; Mayer, R. J., Depletion of 26S Proteasomes in Mouse Brain Neurons Causes Neurodegeneration and Lewy-Like Inclusions Resembling Human Pale Bodies. The Journal of Neuroscience 2008, 28 (33), 8189.

70. Bukhatwa, S.; Zeng, B.-Y.; Rose, S.; Jenner, P., A comparison of changes in proteasomal subunit expression in the substantia nigra in Parkinson’s disease, multiple system atrophy and progressive supranuclear palsy. Brain Research 2010, 1326, 174–183.

71. Savolainen, M. H.; Albert, K.; Airavaara, M.; Myöhänen, T. T., Nigral injection of a proteasomal inhibitor, lactacystin, induces widespread glial cell activation and shows various phenotypes of Parkinson’s disease in young and adult mouse. Exp Brain Res 2017, 235 (7), 2189–2202.

72. Magrinelli, F.; Tesson, C.; Angelova, P. R.; Rodriguez, J. A.; Scardamaglia, A.; O’Callaghan, B.; Lowe, S. A.; Salazar-Villacorta, A.; Chung, B. H.-Y.; Jaconelli, M.; Vona, B.; Esteras, N.; Mammana, A.; Shimazu, J.; Kwong, A. K.-Y.; Courtin, T.; Alavi, S.; Maroofian, R.; Nirujogi, R.; Severino, M.; Monfrini, E.; Rocca, C.; Lewis, P. A.; Efthymiou, S.; Buchert, R.; Sofan, L.; Lis, P.; Pinon, C.; Breedveld, G. J.; Chui, M. M.-C.; Murphy, D.; Pitz, V.; Makarious, M. B.; Baiardi, S.; Volin, M.; Cassar, M.; Hassan, B. A.; Iftikhar, S.; Bauer, P.; Tinazzi, M.; Svetel, M.; Samanci, B.; Hanağası, H. A.; Bilgiç, B.; Cavallieri, F.; Santangelo, M.; Obeso, J. A.; Kurtis, M. M.; Cogan, G.; Kiziltan, G.; Gül-Demirkale, T.; Tireli, H.; Yüksel, G. A.; Yalçın-Cakmakli, G.; Elibol, B.; Barišić, N.; Ng, E. W.-S.; Fan, S.-S.; Hershkovitz, T.; Weiss, K.; Alvi, J. R.; Sultan, T.; Alkhawaja, I. A.; Froukh, T.; Alrukban, H. A. E.; Anjum, M. N.; Saeed, A.; Cheema, H. A.; Fauth, C.; Schatz, U. A.; Zöggeler, T.; Zech, M.; Stals, K.; Varghese, V.; Gandhi, S.; Blauwendraat, C.; Hardy, J. A.; Di Fonzo, A.; Bonifati, V.; Haack, T. B.; Bertoli-Avella, A. M.; Lesage, S.; Başak, A. N.; Steinfeld, R.; Parchi, P.; Jepson, J. E. C.; Alessi, D. R.; Di Fonzo, A.; Brice, A.; Steller, H.; Abramov, A. Y.; Bhatia, K. P.; Houlden, H.; Group, P. S., Variants in the proteasome regulator PSMF1 cause a phenotypic spectrum from parkinsonism to perinatal lethality. Nature Communications 2026.

73. Sullivan, P. G.; Dragicevic, N. B.; Deng, J.-H.; Bai, Y.; Dimayuga, E.; Ding, Q.; Chen, Q.; Bruce-Keller, A. J.; Keller, J. N., Proteasome Inhibition Alters Neural Mitochondrial Homeostasis and Mitochondria Turnover *. Journal of Biological Chemistry 2004, 279 (20), 20699–20707.

74. Tsakiri, E. N.; Gumeni, S.; Vougas, K.; Pendin, D.; Papassideri, I.; Daga, A.; Gorgoulis, V.; Juhász, G.; Scorrano, L.; Trougakos, I. P., Proteasome dysfunction induces excessive proteome instability and loss of mitostasis that can be mitigated by enhancing mitochondrial fusion or autophagy. Autophagy 2019, 15 (10), 1757–1773.

75. Choi, S. J.; Frank, A.; Ben-Johny, M.; Tang, G.; Bendig, J.; Fanourakis, S.; Hobson, B. D.; Ramalingam, N.; Dettmer, U.; Kanter, E.; Sulzer, D.; Mosharov, E. V., α-Synuclein expression is required for somatodendritic dopamine release and immediate early gene induction. Science Advances 2026, 12 (5), eady6978.

76. Geibl, F. F.; Henrich, M. T.; Xie, Z.; Zampese, E.; Ueda, J.; Tkatch, T.; Wokosin, D. L.; Nasiri, E.; Grotmann, C. A.; Dawson, V. L.; Dawson, T. M.; Chandel, N. S.; Oertel, W. H.; Surmeier, D. J., α-Synuclein pathology disrupts mitochondrial function in dopaminergic and cholinergic neurons at-risk in Parkinson’s disease. Molecular Neurodegeneration 2024, 19 (1), 69.

77. Ekstrand, M. I.; Terzioglu, M.; Galter, D.; Zhu, S.; Hofstetter, C.; Lindqvist, E.; Thams, S.; Bergstrand, A.; Hansson, F. S.; Trifunovic, A.; Hoffer, B.; Cullheim, S.; Mohammed, A. H.; Olson, L.; Larsson, N.-G., Progressive parkinsonism in mice with respiratory-chain-deficient dopamine neurons. Proceedings of the National Academy of Sciences 2007, 104 (4), 1325–1330.

78. Stevens, D. A.; Lee, Y.; Kang, H. C.; Lee, B. D.; Lee, Y.-I.; Bower, A.; Jiang, H.; Kang, S.-U.; Andrabi, S. A.; Dawson, V. L.; Shin, J.-H.; Dawson, T. M., Parkin loss leads to PARIS-dependent declines in mitochondrial mass and respiration. Proceedings of the National Academy of Sciences 2015, 112 (37), 11696–11701.

79. Cherra, S. J.; Steer, E.; Gusdon, A. M.; Kiselyov, K.; Chu, C. T., Mutant LRRK2 Elicits Calcium Imbalance and Depletion of Dendritic Mitochondria in Neurons. The American Journal of Pathology 2013, 182 (2), 474–484.

80. Burchell, V. S.; Nelson, D. E.; Sanchez-Martinez, A.; Delgado-Camprubi, M.; Ivatt, R. M.; Pogson, J. H.; Randle, S. J.; Wray, S.; Lewis, P. A.; Houlden, H.; Abramov, A. Y.; Hardy, J.; Wood, N. W.; Whitworth, A. J.; Laman, H.; Plun-Favreau, H., The Parkinson’s disease–linked proteins Fbxo7 and Parkin interact to mediate mitophagy. Nature Neuroscience 2013, 16 (9), 1257–1265.

81. Al Rawi, S.; Simpson, L.; Agnarsdóttir, G.; McDonald, N. Q.; Chernuha, V.; Elpeleg, O.; Zeviani, M.; Barker, R. A.; Spiegel, R.; Laman, H., Study of an FBXO7 patient mutation reveals Fbxo7 and PI31 co-regulate proteasomes and mitochondria. The FEBS Journal 2024, 291 (12), 2565–2589.

82. Liang, C. L.; Wang, T. T.; Luby-Phelps, K.; German, D. C., Mitochondria mass is low in mouse substantia nigra dopamine neurons: implications for Parkinson’s disease. Exp Neurol 2007, 203 (2), 370–80.

83. McWilliams, T. G.; Prescott, A. R.; Montava-Garriga, L.; Ball, G.; Singh, F.; Barini, E.; Muqit, M. M. K.; Brooks, S. P.; Ganley, I. G., Basal Mitophagy Occurs Independently of PINK1 in Mouse Tissues of High Metabolic Demand. Cell Metabolism 2018, 27 (2), 439–449.e5.

84. Guzman, J. N.; Ilijic, E.; Yang, B.; Sanchez-Padilla, J.; Wokosin, D.; Galtieri, D.; Kondapalli, J.; Schumacker, P. T.; Surmeier, D. J., Systemic isradipine treatment diminishes calcium-dependent mitochondrial oxidant stress. The Journal of Clinical Investigation 2018, 128 (6), 2266–2280.

85. Yokota, M.; Kakuta, S.; Shiga, T.; Ishikawa, K. I.; Okano, H.; Hattori, N.; Akamatsu, W.; Koike, M., Establishment of an in vitro model for analyzing mitochondrial ultrastructure in PRKN-mutated patient iPSC-derived dopaminergic neurons. Mol Brain 2021, 14 (1), 58.

86. Rosencrans, W. M.; Lee, R. W.; McGraw, L.; Horsburgh, I.; Wang, T.-Y.; Quan, B.; Huynh, D.; Johnston, J. A.; Chan, D. C.; Chou, T.-F., Putative PINK1/Parkin activators lower the threshold for mitophagy by sensitizing cells to mitochondrial stress. Science Advances 11 (35), eady0240.

87. Sarchione, A.; Marchand, A.; Taymans, J.-M.; Chartier-Harlin, M.-C., Alpha-Synuclein and Lipids: The Elephant in the Room? Cells 2021, 10 (9), 2452.

88. Ellis, C. E.; Murphy, E. J.; Mitchell, D. C.; Golovko, M. Y.; Scaglia, F.; Barceló-Coblijn, G. C.; Nussbaum, R. L., Mitochondrial lipid abnormality and electron transport chain impairment in mice lacking alpha-synuclein. Mol Cell Biol 2005, 25 (22), 10190–201.

89. Rappold, P. M.; Cui, M.; Grima, J. C.; Fan, R. Z.; de Mesy-Bentley, K. L.; Chen, L.; Zhuang, X.; Bowers, W. J.; Tieu, K., Drp1 inhibition attenuates neurotoxicity and dopamine release deficits in vivo. Nature Communications 2014, 5 (1), 5244.

90. Brown, H. J.; Fan, R. Z.; Bell, R.; Salehe, S. S.; Martínez, C. M.; Lai, Y.; Tieu, K., Imbalanced mitochondrial dynamics in human PD and α-synuclein mouse brains. Neurobiology of Disease 2025, 212, 106976.

91. Bordt, E. A.; Zhang, N.; Waddell, J.; Polster, B. M., The Non-Specific Drp1 Inhibitor Mdivi-1 Has Modest Biochemical Antioxidant Activity. Antioxidants (Basel) 2022, 11 (3).

92. Bordt, E. A.; Clerc, P.; Roelofs, B. A.; Saladino, A. J.; Tretter, L.; Adam-Vizi, V.; Cherok, E.; Khalil, A.; Yadava, N.; Ge, S. X.; Francis, T. C.; Kennedy, N. W.; Picton, L. K.; Kumar, T.; Uppuluri, S.; Miller, A. M.; Itoh, K.; Karbowski, M.; Sesaki, H.; Hill, R. B.; Polster, B. M., The Putative Drp1 Inhibitor mdivi-1 Is a Reversible Mitochondrial Complex I Inhibitor that Modulates Reactive Oxygen Species. Dev Cell 2017, 40 (6), 583–594.e6.

93. Kriks, S.; Shim, J. W.; Piao, J.; Ganat, Y. M.; Wakeman, D. R.; Xie, Z.; Carrillo-Reid, L.; Auyeung, G.; Antonacci, C.; Buch, A.; Yang, L.; Beal, M. F.; Surmeier, D. J.; Kordower, J. H.; Tabar, V.; Studer, L., Dopamine neurons derived from human ES cells efficiently engraft in animal models of Parkinson’s disease. Nature 2011, 480 (7378), 547–51.

94. Jenkins, B.; Ronis, M.; Koulman, A., LC–MS Lipidomics: Exploiting a Simple High-Throughput Method for the Comprehensive Extraction of Lipids in a Ruminant Fat Dose-Response Study. Metabolites 2020, 10 (7), 296.

95. Leestemaker, Y.; de Jong, A.; Witting, K. F.; Penning, R.; Schuurman, K.; Rodenko, B.; Zaal, E. A.; van de Kooij, B.; Laufer, S.; Heck, A. J. R.; Borst, J.; Scheper, W.; Berkers, C. R.; Ovaa, H., Proteasome Activation by Small Molecules. Cell Chemical Biology 2017, 24 (6), 725–736.e7.

96. Jerber, J.; Haldane, J.; Steer, J.; Pearce, D.; Patel, M., Dissociation of neuronal culture to single cells for scRNA-seq (10x Genomics) v1. 2020.

97. Alles, J.; Karaiskos, N.; Praktiknjo, S. D.; Grosswendt, S.; Wahle, P.; Ruffault, P.-L.; Ayoub, S.; Schreyer, L.; Boltengagen, A.; Birchmeier, C.; Zinzen, R.; Kocks, C.; Rajewsky, N., Cell fixation and preservation for droplet-based single-cell transcriptomics. BMC Biology 2017, 15 (1), 44.

