## Supplementary Figures for "Pathological levels of α-synuclein increase basal mitophagy in Parkinson’s dopamine neurons"

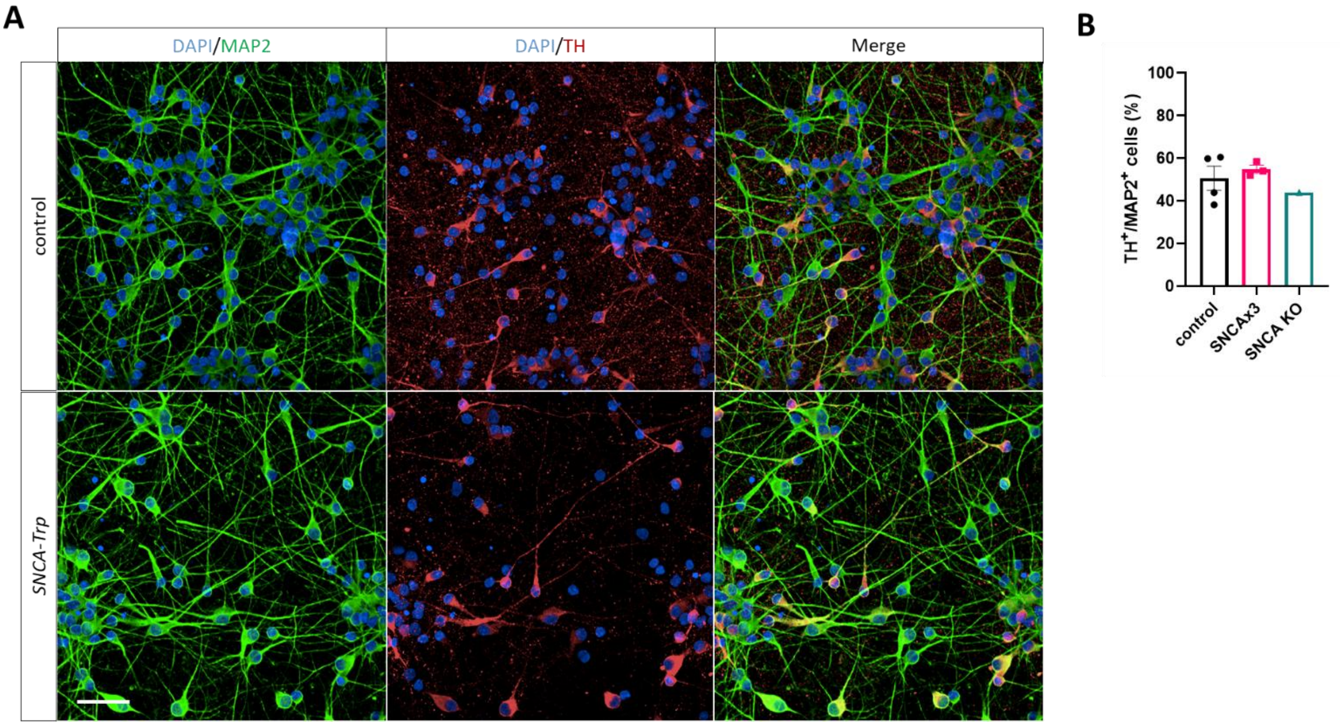


**Supplementary Fig. 1. A)** Representative images of control and *SNCA-Trp* MAP2-positive TH-positive iPSC-DANs at D41 of differentiation. Scale bar = 50 µm. **B)** Quantification of the percentage of $TH^+$ $MAP2^+$ cells showing no significant difference between genotypes. Data represent the mean ± SEM of 3-4 lines of iPSC-DANs per genotype averaged for three independent differentiations.

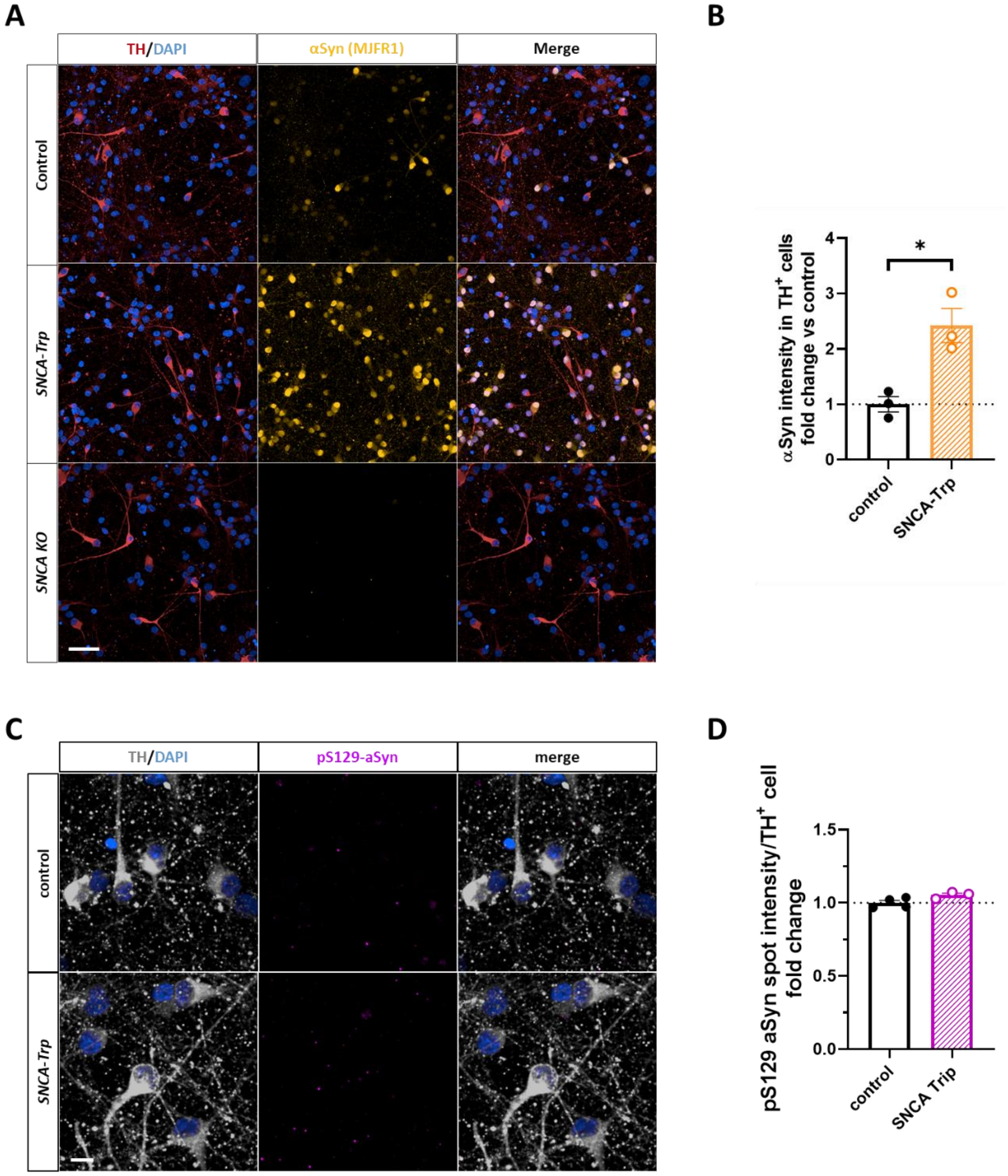


**Supplementary Fig. 2. A)** Representative images showing a 2-3 fold increase in α-synuclein expression in *SNCA-Trp* TH-positive iPSC-DANs compared to controls at D40. The specificity of the total α-synuclein (MJFR1) antibody was confirmed using *SNCA KO* iPSC-DANs. Scale bar = 50 µm. **B)** Quantification of α-synuclein immunofluorescence as depicted in **(A)**. Data represent the mean ± SEM of 3 lines of iPSC-DANs per genotype averaged over two independent differentiations. *P*-values were calculated using unpaired, two-tailed Student's *t*-test with Welch's correction. **C)** Representative images of pS129-aSyn immunofluorescence in control and SNCA-Trp iPSC-DANs. Scale bar = 10 µm. **D)** No increase in α-synuclein pS129 intensity was observed in $TH^+$ iPSC-DANs. *, $P < 0.05$.

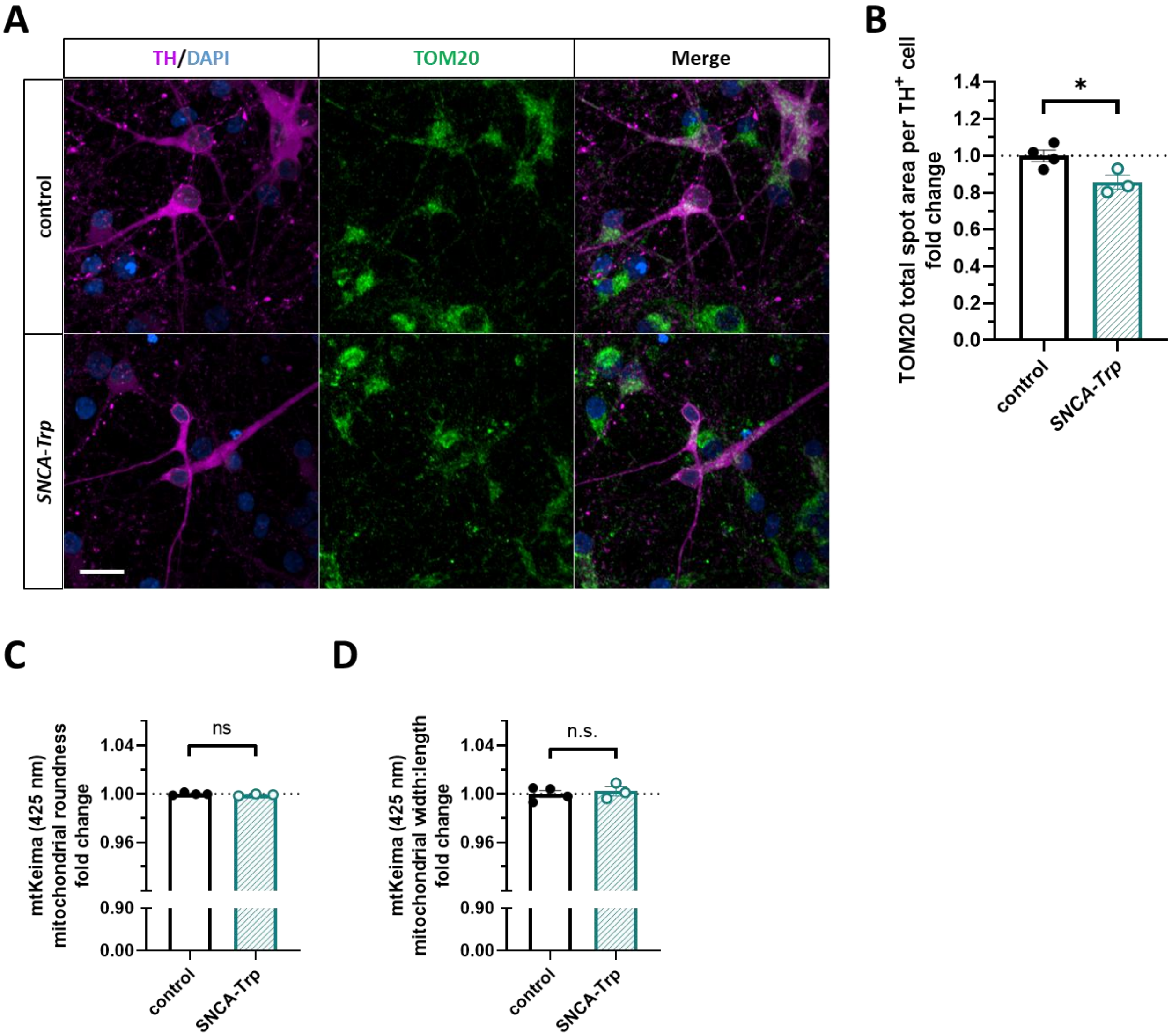


**Supplementary Fig. 3. A)** Representative images of TH⁺ iPSC-DANs stained for mitochondria (TOM20). Scale bar = 20 µm. **B**) Quantification of (**A**) showing decreased TOM20 total spot area in TH⁺ *SNCA-Trp* iPSC-DANs vs control. Data represent the mean ± SEM of 3-4 lines of iPSC-DANs per genotype averaged for three independent differentiations. *P*-values were calculated using unpaired, two-tailed Student's *t*-test with Welch's correction. **C-D**) Quantification of green healthy mt-keima mitochondria related to Fig. 1 showing no significant change in (**C**) roundness and (**D**) width:length ratio in *SNCA-Trp* vs control iPSC-DANs. For **C-D**) data represent the mean ± SEM of 3-4 lines of iPSC-DANs per genotype averaged over four independent differentiations. *, $P < 0.05$.

**A**

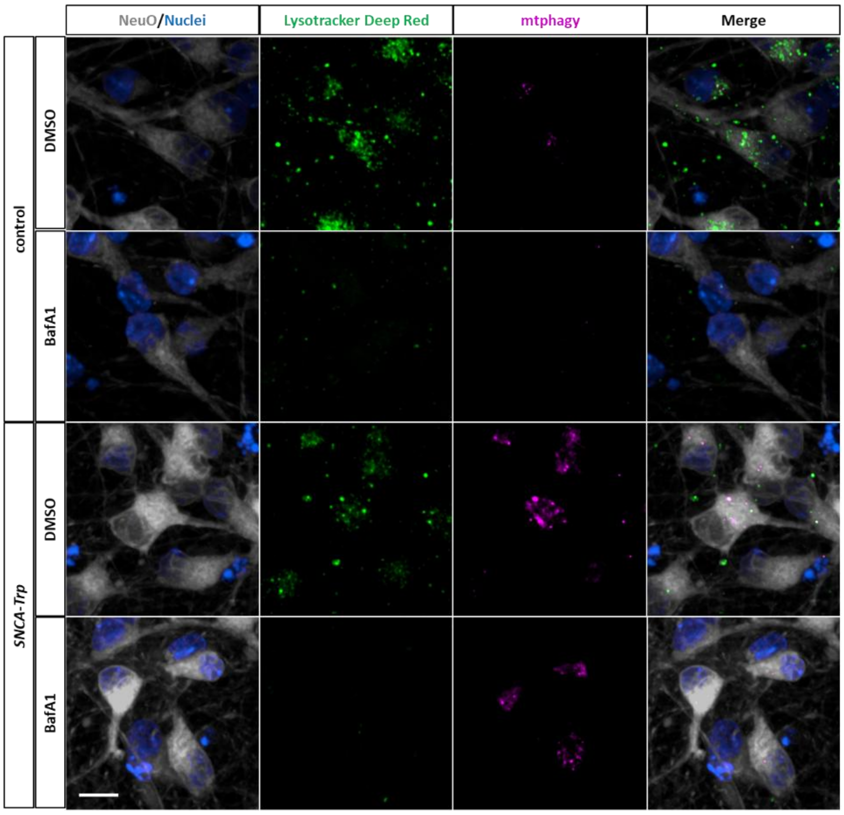


**B**

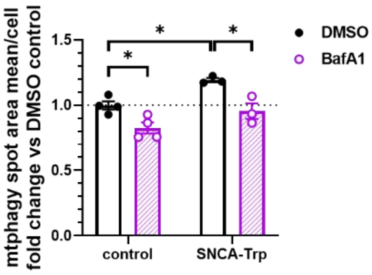


**Supplementary Fig. 4. mtPhagy mitophagy reporter shows larger mitolysosomes in SNCA-Trp iPSC-DANs. A)** Representative images of iPSC-DANs treated with mtPhagy dye (50 nM, 48 h, magenta) and co-stained with Lysotracker Deep Red (100 nM, 3 h, green). Bafilomycin A1 (BafA1, 500 nM) was added 3 h before imaging. **B)** Quantification of mtPhagy+/Lysotracker Deep Red+ puncta mean spot area. Data represent the mean ± SEM of 3-4 lines of iPSC-DANs per genotype. *P*-values were calculated using two-way ANOVA with Tukey's multiple comparisons test. *, $P < 0.05$; **, $P < 0.01$; *** $P < 0.001$.

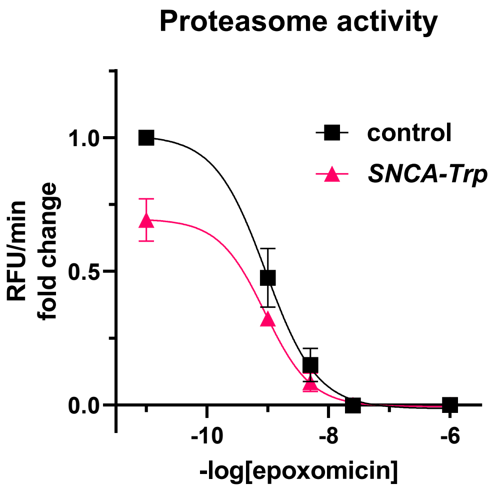


**Supplementary Fig. 5.** Dose-response curves of epoxomicin inhibiting chymotrypsin-like (β5) proteasome activity in control and *SNCA-Trp* iPSC-DANs. For control: $IC_{50}$ = 0.93 nM, 95% CI: 0.23 – 1.7 nM; *SNCA-Trp* $IC_{50}$ = 0.90 nM, 95% CI: 0.25 – 1.6 nM. Data represent the mean ± SEM from 3 lines of iPSC-DANs per genotype averaged over two independent differentiations.

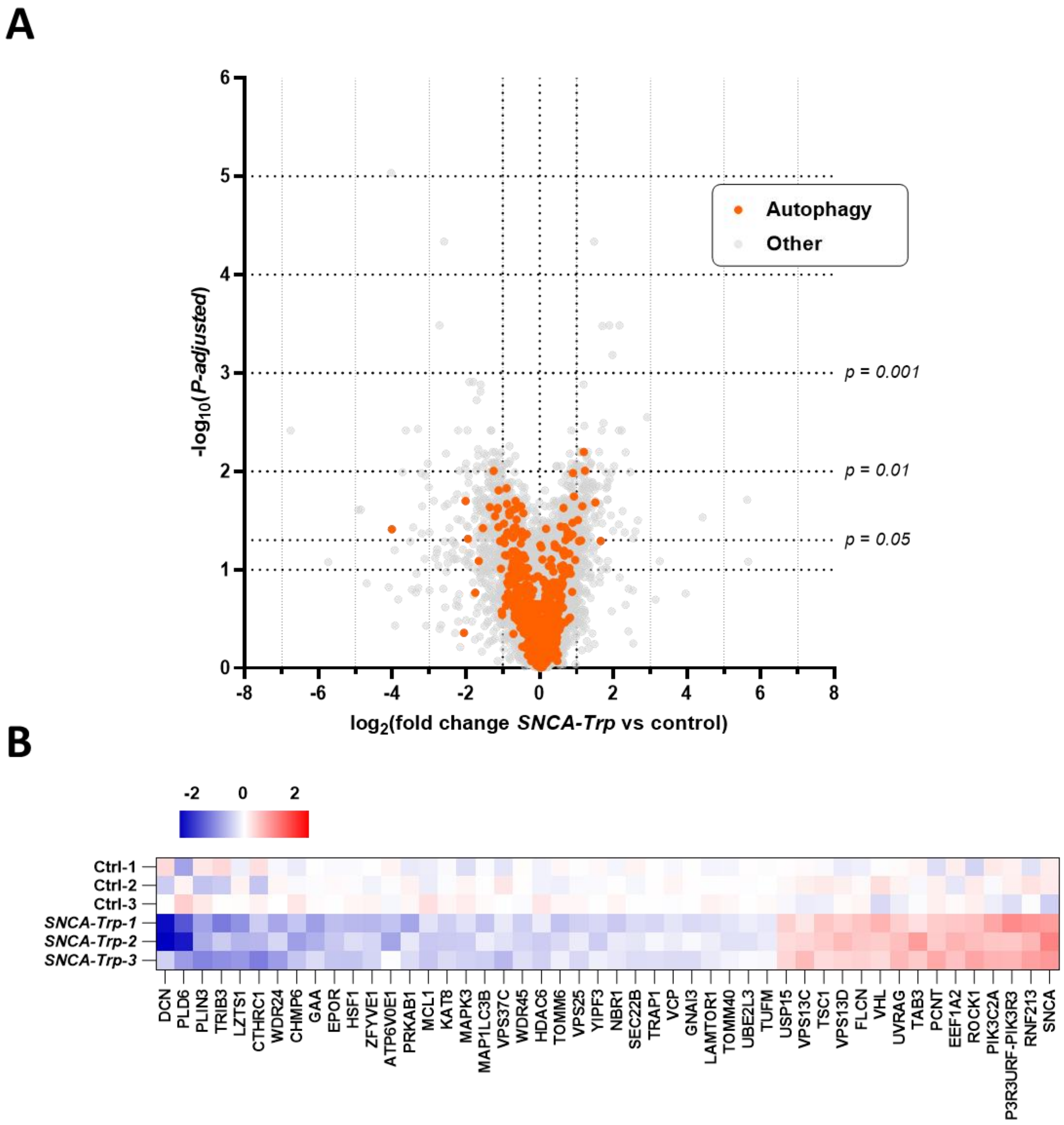


**Supplementary Figure 6. scRNA-seq shows *SNCA-Trp* iPSC-DANs have downregulated expression of autophagy genes. A**) Volcano plot depicting differentially expressed autophagy-related genes (orange) from the DGEA. **B**) Heat map displaying $log_2$ transformed differential expression per iPSC-derived neuronal cell line for significantly (*P*-adjusted < 0.05) differentially expressed autophagy genes. For **A**), data represent the mean $log_2$ fold change ratio from the DGEA for 3 controls and 3 *SNCA-Trp* lines of iPSC-DANs with Benjamini-Hochberg adjusted *P*-value (FDR < 0.05).

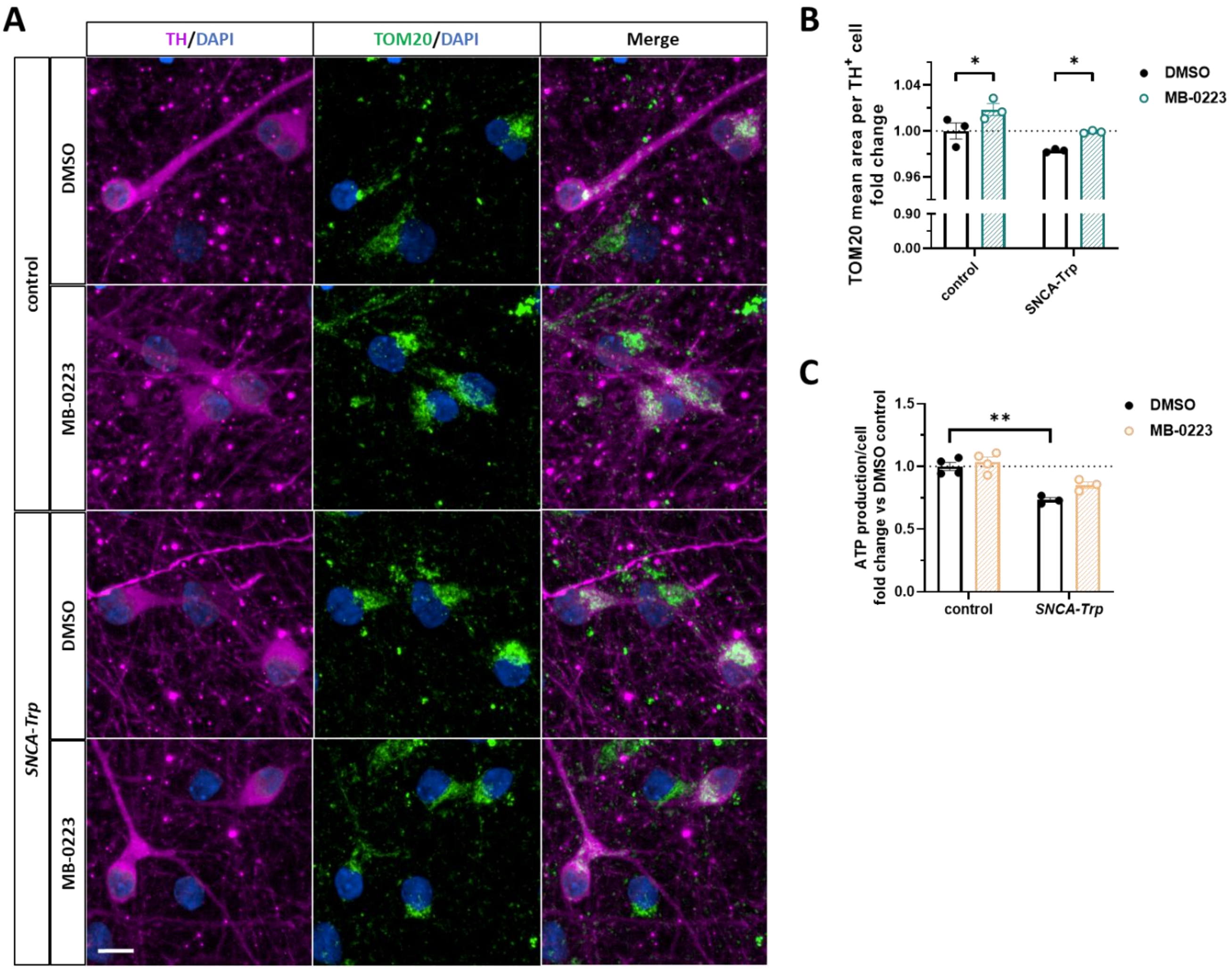


**Supplementary Fig.7. DRP1 inhibitor MB-0223 increases mitochondrial size in TH⁺ iPSC-DANs and restores ATP levels. A)** Representative images of TH⁺ dopaminergic neurons showing increased TOM20 mean spot area upon treatment with MB-0223 (3 µM, 48 h). Scale bar = 10 µm. **B**) Quantification of TOM20 immunofluorescence. Data represent the mean ± SEM of 3-4 lines of iPSC-DANs per genotype averaged for three independent differentiations. **C**) ATP production as measured by luminescence is significantly decreased in *SNCA-Trp* iPSC-DANs and reversed by MB-0223 (3 µM, 48 h). *P*-values were calculated using two-way ANOVA with Tukey's multiple comparisons test. *, $P < 0.05$; **, $P < 0.01$.

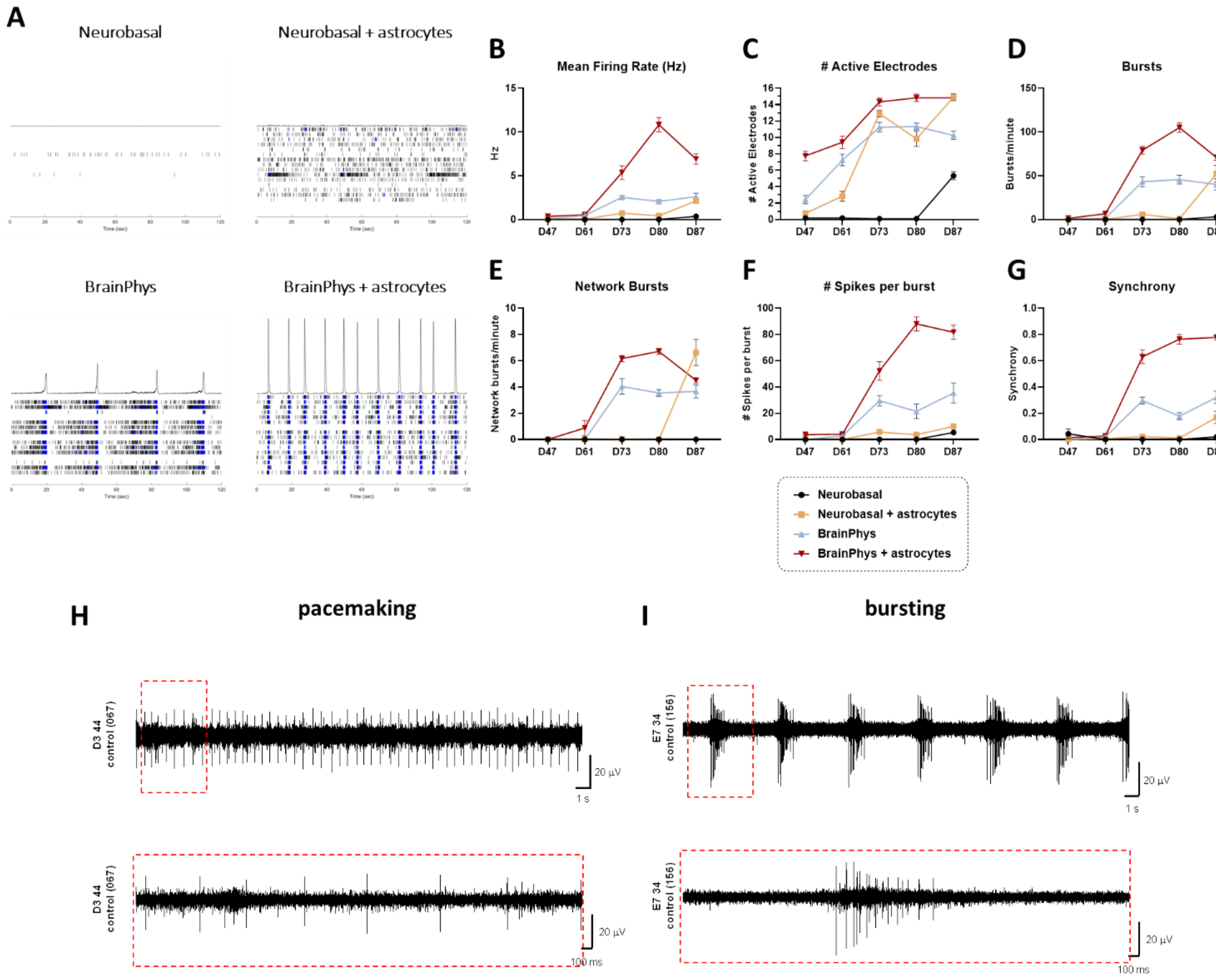


**Supplementary Fig. 8. BrainPhys and rat astrocytes increase synchronised neuronal activity in iPSC-DANs. A)** Representative spike raster plots depicting increased firing activity in BrainPhys alone or BrainPhys + astrocytes in iPSC-DANs from a healthy control (D101). **B-G)** Quantification of firing activity **(B)**, number of active electrodes **(C)**, bursts per minute **(D)**, network bursts per minute **(E)**, number of spikes per burst **(F)**, and synchrony **(G)**. Data represent the mean ± SEM of twelve technical replicates for 1 differentiation in a control line of iPSC-DANs. **G,H**) Single electrode field potentials showcasing dopaminergic pacemaking (**G**) or bursting (**H**) activity in a control line of iPSC-DANs.

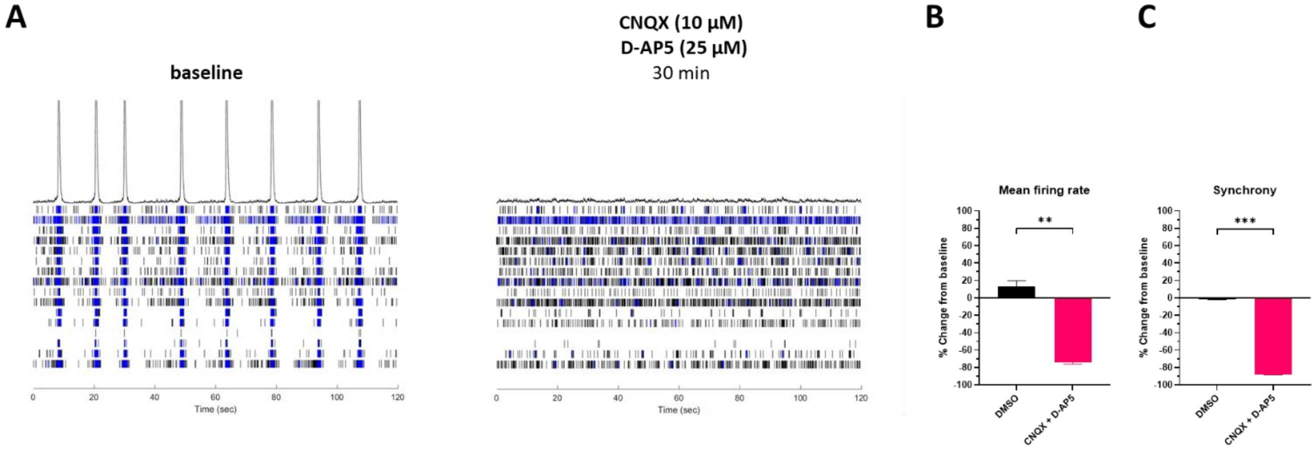


**Supplementary Fig. 9. Synchronised network bursting activity in iPSC-DANs is dependent on AMPA and NMDA receptors.** **A)** Representative spike raster plots showing baseline synchronised network bursting activity in iPSC-DANs from a healthy control, which is abolished by blocking AMPA and NMDA receptors with CNQX (10 µM) and D-AP5 (25 µM), respectively. **B+C)** Quantification shows significant loss of firing activity **(B)** and synchrony **(C)**. Data represent the mean ± SEM of four technical replicates for 1 differentiation in a control line of iPSC-DANs. *, $P < 0.05$; **, $P < 0.01$; *** $P < 0.001$.

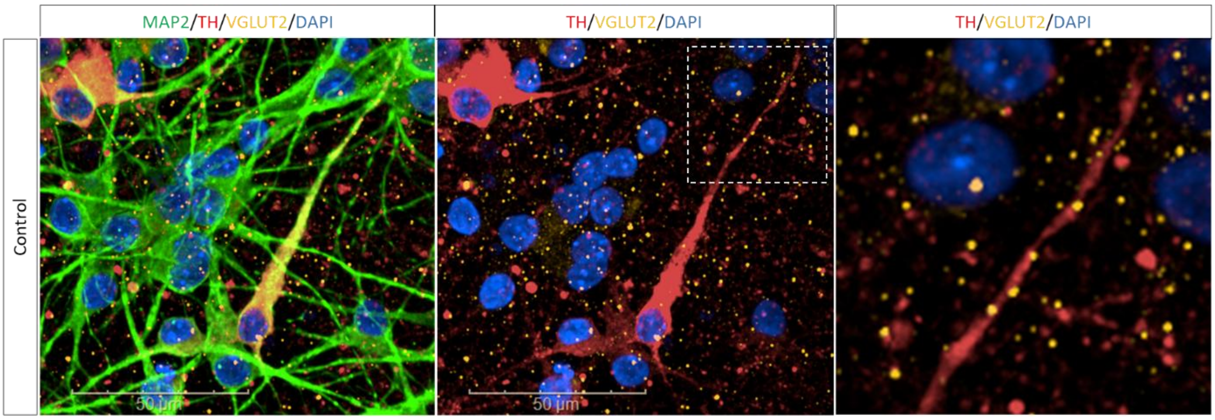


**Supplementary Fig 10. TH-positive iPSC-DANs co-express VGLUT2 in soma and axons.** Immunostaining showing VGLUT2$^{+}$ puncta in TH$^{+}$ dopamine neurons from a healthy control at D70. Dopamine neurons were co-cultured with rat astrocytes.
